# TGFβ1 and RGD Cooperatively Regulate SMAD 2/3 Mediated Oncogenic Effects in Prostate Cancer Cells in Bioorthogonally Constructed Hydrogels

**DOI:** 10.1101/2024.12.09.627597

**Authors:** Mugdha Pol, Hanyuan Gao, Joseph M Fox, Xinqiao Jia

**Author notes:** Corresponding author: Dr. Xinqiao Jia, Ammon-Pinizzotto Biopharmaceutical Innovation Center, 590 Avenue 1743, Newark, DE 19713, USA.

## Abstract

To recapitulate prostate cancer metastasis, DU145 cells were cultured in a hyaluronic acid-based, bioorthogonally constructed, protease-degradable hydrogel. In the presence of covalently conjugated integrin-binding peptide (G<u>RGD</u>SP), DU145 cells formed tumoroids and exhibited small protrusions. Upon addition of soluble transforming growth factor beta 1 (TGFβ1), cells underwent morphological changes to form extended interconnected cellular networks. Contrarily, in RGD-free hydrogels, cells maintained spherical structures even in the presence of TGFβ1. In RGD-conjugated hydrogels, TGFβ1 induced nuclear localization of SMAD2/3, upregulating a wide range of TGFβ1 target genes and proteins. Prolonged exposure to TGFβ1 led to matrix remodeling and induced epithelial-to-mesenchymal transition in DU145 cells, with loss of epithelial markers and gain of mesenchymal markers. TGFβRI/ALK5 inhibitor SB-431542 attenuated TGFβ1-induced morphological changes, abrogated nuclear localization of SMAD2/3, and restored the expression of key epithelial markers. Our findings highlight the cooperative role of TGFβ1 signaling and integrin binding peptide in the acquisition of aggressive phenotype and promoting tumor progression.

**TEASER:** Physiologically relevant 3D cell culture platforms enabled mechanistic investigation of growth factor signaling related to prostate cancer metastasis.

## INTRODUCTION

Prostate cancer (PCa) is the most diagnosed malignancy and the second leading cause of cancer- related deaths in men worldwide^1^. Nearly 80% of men are diagnosed with organ-confined malignancy, 15% with local tissue or pelvic lymph node invasion, and 5% with distant metastases^2^. While life expectancy for men with localized disease can be as high as 90% over 10 years, prognosis with advanced disease drops down to only 30% at 5 years, accompanied by poor quality of life^3^. The mechanisms of prostate cancer initiation and progression are poorly understood, as prostate cancer is characterized by its heterogenicity, with multiple tumor foci being synchronously present in the organ of origin^4, 5^. It is well-established that tumors exhibit a predilection for metastasis to specific organs. Extensive experimental and clinical data has confirmed the “seed and soil” hypothesis, that organ preferences of tumor metastasis are due to favorable interactions between cancer cell’s innate features (seed) and the availability of a conducive organ microenvironment (soil)^6, 7^.

During tumorigenesis, communication between different stromal and cancer cells is controlled by cytokines, which act in an autocrine, paracrine, or juxtacrine manner to remodel the tumor microenvironment^8^. Transforming growth factor-beta (TGFβ), the most pleiotropic known cytokine, plays a prominent role in cancer progression. The transforming growth factor β family is encoded by 33 genes and includes three isoforms, TGFβ1, TGFβ2, and TGFβ3, which are 75% homologous with analogous signaling events but exhibiting variable expression based on the cell type^9, 10^. TGFβ1 is the most dysregulated isoform in cancer progression and is predominantly expressed in connective tissue, hematopoietic, and endothelial cells^11^. After TGFβ1 binds to the type 2 receptor (TGFBR2), the type 1 receptor/activin receptor-like kinase (TGFBR1/ALK5) is recruited and phosphorylated, resulting in downstream signaling activation. The downstream signaling proceeds either through SMAD-independent non-canonical signaling or SMAD-mediated canonical signaling. The canonical signaling pathway is initiated when the carboxy-terminal serine residue of SMAD2/3 is phosphorylated. Phosphorylation of SMAD2/3 induces its oligomerization with SMAD4, which promotes nuclear translocation of SMAD2/3/4 complex. SMAD complex nuclear localization induces transcriptional activation or repression via its interaction with transcription cofactors, thereby inducing morphological and phenotypic changes^12^. TGFβ1 is known as a tumor suppressor by inducing cytostatic effect and as a tumor promoter leading to cancer metastasis^13^. Prolonged exposure to TGFβ1 induced epithelial-to-mesenchymal transition (EMT) in epithelial cells, which triggers a switch from the non-motile epithelial to a motile mesenchymal state^14–17^. The transformed cells are aggressive, invasive, and resistant to apoptosis, senescence, immune surveillance, and chemotherapy^18^.

A key challenge in clinical management and therapeutic intervention of PCa is the lack of relevant preclinical models that accurately recapitulate the complex 3D microenvironment and tumor heterogenicity. Although genetically engineered mouse models and patient-derived xenografts are widely used and have significantly advanced the field, these models do not permit in-depth investigations into how the complex cellular and extracellular microenvironments impact cell fate decisions due to the limited control of the tumor microenvironment^19, 20^. This is further complicated by the inherent animal-to-animal variation, the high cost, and the heavy workload^21^. Over the past few decades, biomaterials-based engineered tumor models have gained recognition in academia and pharmaceutical industries. These reductionist platforms allow researchers to parse the complex *in vivo* tumor biology to examine features of the extracellular microenvironment and their influence on cell behaviors. By analyzing cell growth, cell-cell, cell-extracellular matrix (ECM) interaction, homeostasis, invasion, EMT, and eventually metastasis in the engineered models, new anticancer therapies have been developed^21–23^.

Previously, we hreported a hyaluronic acid (HA)-based bioorthogonal hydrogel platform that promotes the formation of 3D tumoroids from dispersed single cells and permits user-directed matrix remodeling in a time-delayed manner to induce cell dissemination from the spheroids and invasion into the synthetic matrix. DU145 cells, a brain metastasized prostate cancer cell line, were encapsulated in the HA gel via the slow (second-order rate constant, *k_2_* ∼ 10^-2^-10^0^ M^-1^s^-1^) tetrazine (Tz)-norbornene (Nb) cycloaddition reaction. Seven days later, fibronectin-derived, integrin-binding peptide (G<u>RGD</u>SP) was conjugated to the matrix by employing the rapid (*k_2_* ∼ 10^5^-10^6^ M^-1^s^-1^) and highly efficient tetrazine ligation with *trans*-cyclooctene (TCO). In response, DU145 cells underwent EMT and developed cortactin-positive invadopodia around the tumoroids^24, 25^. While the synthetic ECM served as a framework for PCa cells to adhere, expand, and migrate, the potent regulatory roles of TGFβ1 in cancer progression were not evaluated. Herein, one day after cell encapsulation, TCO-tagged RGD peptide was supplemented in the cell culture media to allow for instantaneous immobilization of the cell binding motif at the gel-liquid interface. On day 3, soluble TGFβ1 was introduced, and cultures were maintained with TGFβ1 for 21 days (Fig. 1A). Using the modular 3D culture system, we demonstrate that matrix adhesiveness primes the resident PCa cells to respond to TGFβ1, SMAD2/3 nuclear localization induces transcriptional changes in TGFβ1 target genes, and activation of TGFβ signaling leads to EMT. Control experiments with a TGFβRI/ALK5 inhibitor (SB-431542) further corroborated our findings. Overall, this study demonstrates the cooperative effects of soluble (TGFβ1) and insoluble (RGD) ECM signals in promoting cancer metastasis.

**Fig. 1.**
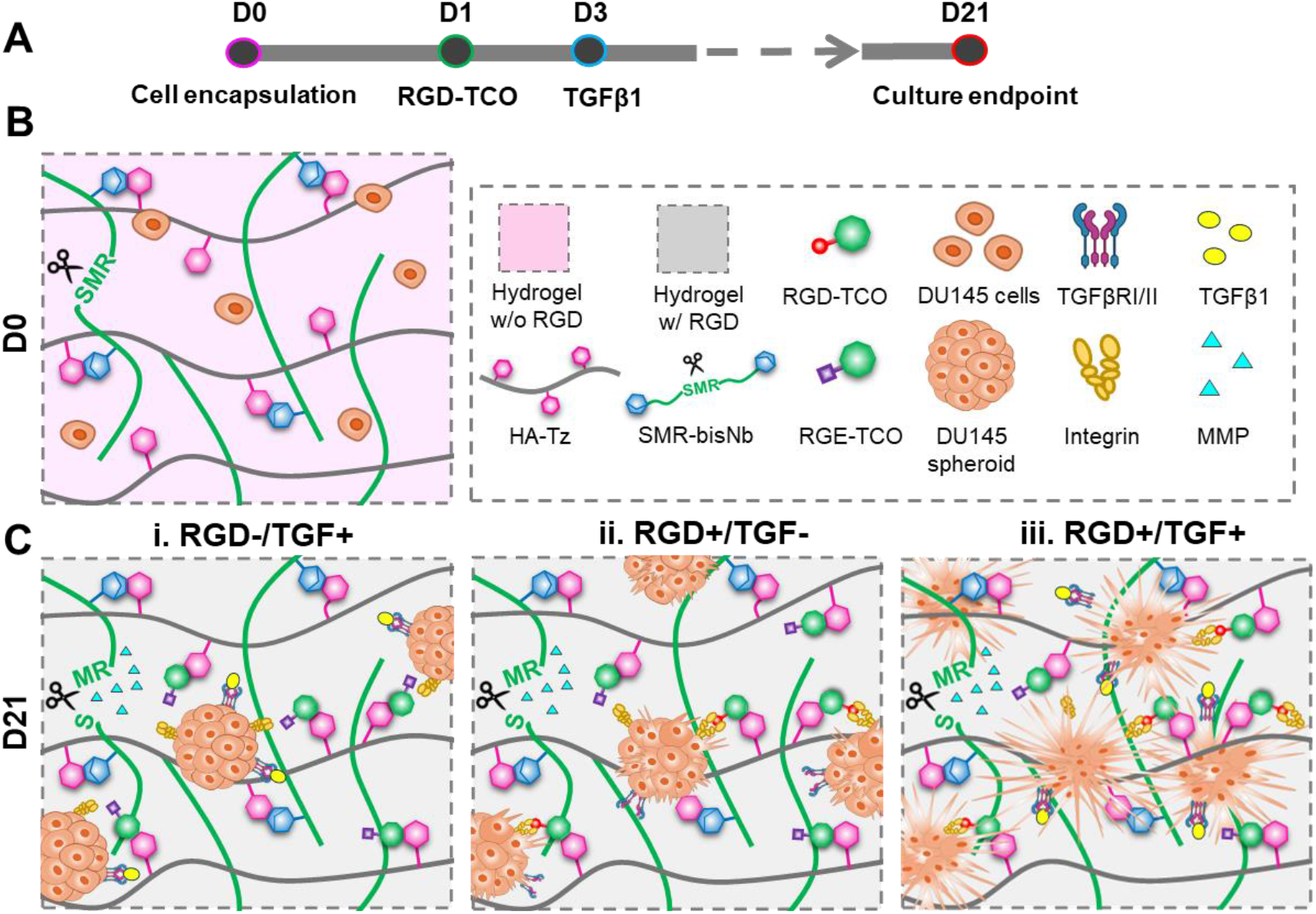
Schematic depiction of TGFβ1-induced EMT in PCa cells cultured in a biomimetic HA-based hydrogel. **(A)** Flow chart showing the experimental timeline. On day 0, DU145 cells were dispersed in a mixture of HA-Tz and SMR-bisNb at a Tz/Nb molar ratio of 2/1. On day 1, TCO-tagged peptide ligands (RGD-TCO) were added to the media to allow the diffusion-controlled Tz-TCO reaction to proceed for 24 h. On day 3, TGFβ1 was introduced to the media at 10 ng/mL. Cultures were maintained in TGFβ1 conditioned media until day 21. **(B)** DU145 single cells cultured in MMP-degradable HA gels without the cell adhesive motif on day 0. **(C)** Three types of cellular constructs developed on day 21. DU145 cells maintained in RGD-free HA gels in the presence of TGFβ1 spontaneously assembled into compact multicellular spheroids (i). In the absence of TGFβ1, multicellular structures developed in RGD- containing, MMP-degradable gels showed small protrusions at the edge of the spheroids (ii). When both RGD and TGFβ1 signals were present, extensive cell dissemination from the spheroids and invasion into the matrix were observed (iii). Figures for hydrogels were created using BioRender.com.

## RESULTS

The HA-based hydrogel platform permits *in situ* modulation of matrix properties during 3D live cell culture^25^. The hydrogel building blocks include tetrazine-modified HA (HA-Tz), TCO- functionalized cell adhesive peptide RGD-TCO or non-adhesive control RGE-TCO, norbornene- functionalized, protease degradable crosslinker SMR-bisNb (VPMS↓MRGG) ^24, 26, 27^, and the scrambled, non-degradable peptide crosslinker, SMV-bisNb (SMVGMRPG)^28^ (Fig. S1-S3). Mixing of HA-Tz and SMR-bisNb at a Tz/Nb molar ratio of 2/1 gave rise to hydrogels with an optimal stiffness required to grow DU145 cells^29^ in 3D. One day later, RGD was homogeneously conjugated to the matrix via diffusion-controlled bioorthogonal ligation at the gel-liquid interface. Because Tz/TCO ligation is highly efficient and instantaneous, the composition of the TCO reservoir, i.e. the molar ratio of RGD-TCO relative to RGE-TCO, was reflected in the modified hydrogel in 3D, thereby enabling precise tuning of the RGD concentrations in the final hydrogel construct. Cells were maintained in 3D in TGFβ1 conditioned media from days 3 to 21. We evaluated how matrix degradability and adhesiveness mediated cell growth, morphology, phenotype, and TGFβ1 signaling (Fig. 1B).

### RGD-decorated, MMP-degradable matrix is required for TGFβ1 regulation in 3D

To confirm the regulatory effects of TGFβ1 on DU145 cells, a 2D wound scratch assay was performed using GFP-labeled DU145 cells (Fig. 2A). In the presence of TGFβ1, the wound was completely closed in 24 h, in sharp contrast to the partial closure observed in the untreated controls. Compared to the TGFβ1-free controls, TGFβ1 treated DU145 cells were larger and more elongated (Fig. S4), resembling the spindle-like morphology seen in mesenchymal cells, with reduced cell- cell contacts. These results confirm that DU145 cells respond to TGFβ1 on 2D by altering cell morphology and increasing cell migration.

**Fig. 2.**
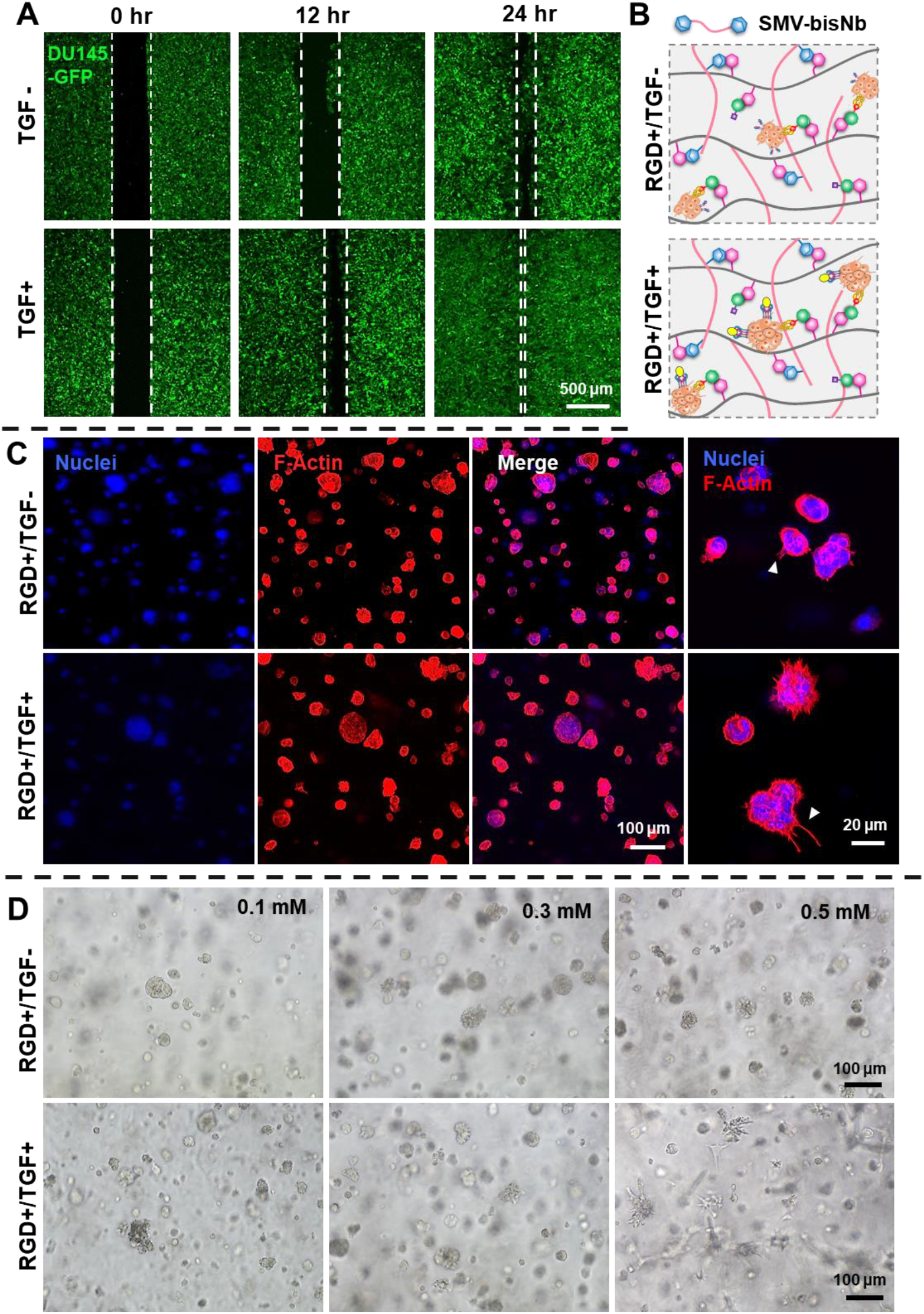
Characterization of TGFβ1-induced immediate responses from DU145 cells cultured on 2D (A) and identification of appropriate matrix parameters for 3D cultures (B-D). **(A)** Representative fluorescent images showing the progression of the scratch/wound closure for GFP-labeled DU145 cells at 0, 12, and 24 h. Scale bar: 500 µm. **(B)** Schematic depiction of PCa cells cultured in HA-gels that were not MMP degradable (crosslinker SMV-bisNb) in the presence of RGD and TGFβ1. (**C)** Representative confocal images of DU145 cells cultured in RGD (1.0 mM)-containing HA gels that were not MMP-degradable for 21 days with or without TGFβ1 (10 ng/mL). Cell nuclei were stained blue by DAPI, and F-actin was stained red by Alexa Fluor^TM^ 568 phalloidin. Arrowheads point to small protrusive structures. Scale bar: 100 µm (left panel) and 20 µm (right panel). **(D)** Bright-field images of cells cultured in MMP-degradable hydrogels with varying amounts of RGD (0.1, 0.3, and 0.5 mM) with or without TGFβ1 (10 ng/mL) for 21 days. Scale bar: 100 µm.

We then screened the biomaterials space to identify appropriate matrix conditions that activate TGFβ signaling and promote cell invasion in 3D. DU145 cells were cultured in hydrogels that were not susceptible to MMP-mediated cleavage using the scrambled peptide crosslinker (SMV- bisNb)^28^. In these experiments, cells were maintained in gels with 1.0 mM RGD, with or without TGFβ1 (Fig. 2B). Individually dispersed single cells proliferated moderately and formed small multicellular spheroids by day 7 irrespective of whether TGFβ1 was present in the media (Fig. 2C, Fig. S5). In both constructs, spheroids appeared larger by day 21, although significant amounts of single cells were still present. All multicellular structures remained small and compact. Small hair- like structures resembling filopodia were seen radiating from the edges of the spheroids in both constructs. Even in the presence of TGFβ1, spheroids did not loosen cell-cell contacts or develop protrusive structures. Overall, in RGD-containing, non-degradable matrices, soluble TGFβ1 could not exert its potent regulatory functions.

After confirming the need for MMP- cleavable crosslinks, we queried the optimal RGD density for maximized activation of TGFβ1 signaling. DU145 cells were maintained in SMR-bisNb- crosslinked HA gels with RGD concentrations ranging from 0.1 to 2.5 mM. In all cases, spheroid formation was observed by day 21, although spheroid size and morphology differed significantly (Fig. 2D). Under RGD+/TFG- conditions, spheroids developed in gels with 0.1 mM RGD were sparse, small, and compact with defined circular borders, while those grown in gels with 0.3 and 0.5 mM RGD were larger, more abundant, and had rugged borders. Adding soluble TGFβ1 did not significantly alter the morphology of the multicellular structures developed in cultures with 0.1 and 0.3 mM RGD but led to the establishment of invadopodia- like protrusive structures around individual cell clusters in cultures with 0.5 mM RGD. In the absence of TGFβ1, irregularly shaped, loose spheroids with small cellular protrusions were detected in gels with 1.0 mM RGD (Fig. 3A: day 21/RGD+/TGF-). Strikingly, when this culture was supplemented with TGFβ1, a dense cellular network with extended cellular processes was developed (Fig. 3A, day 21/RGD+/TGF+). In the absence of TGFβ1 with 2.5 mM RGD, markedly larger spheroids were observed. The structures were morphologically more heterogeneous, with some compact and spherical, others loose and scattered, and others exhibiting cellular extensions radiating from the spheroid body (Fig. S6: day 21). Collectively, to respond to TGFβ1 signaling, PCa cells must be maintained in a protease-degradable matrix with a minimum RGD concentration of 1.0 mM. All subsequent experiments were carried out using SMR-bisNb-crosslinked hydrogels with 1.0 mM RGD. RGD-free hydrogels were included as the controls.

**Fig. 3.**
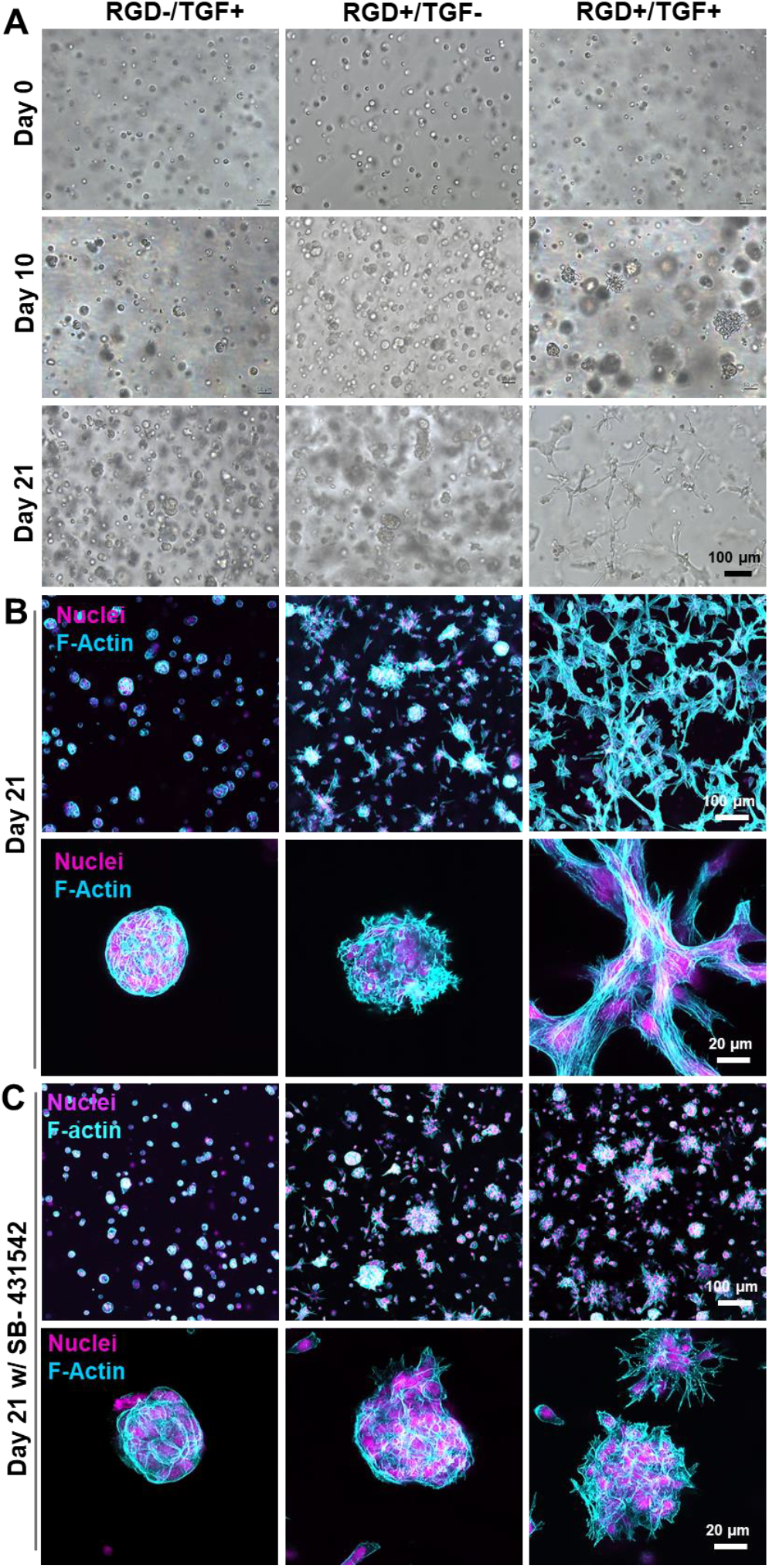
TGFβ1 induces PCa morphogenesis in RGD primed hydrogels. **(A)** Bright-field images of 3D cultures with or without RGD (1.0 mM) and/or TGFβ1 (10 ng/mL) on days 0, 10 and 21. Scale bar: 100 µm. **(B)** Representative confocal images of day 21 cultures fluorescently stained for nuclei (DAPI, magenta) and F-actin (Alexa Fluor^TM^ 568 phalloidin, cyan). Scale bar: 100 µm (top panel), 20 µm (bottom panel). **(C)** Representative confocal images of day 21 cultures maintained with SB-431542 (10 μM, supplemented in cultures from day 3 to day 21) and fluorescently stained for nuclei with DAPI (magenta) and F-actin (cyan). Scale bar: 100 µm (top panel) and 20 µm (bottom panel).

### RGD and TGFβ1 synergistically promote cell invasion in 3D

Homogenously dispersed DU145 single cells underwent clonal expansion to form small spheroids, each containing approximately 4-6 cells by day 3 under all three experimental conditions (Fig. S7: RGD-/TGF+, RGD+/TGF- and RGD+/TGF+). RGD immobilization on day 1 supported the rapid growth of larger spheroids, and by day 5, compact spheroids were detected, measuring on average 50 μm (Fig. S8). Spheroid size continued to increase over time in all three constructs and for RGD-/TGF+ and RGD+/TGF- constructs, cells remained closely packed within the spheroid by day 10. Meanwhile, spheroids in RGD+/TGF+ cultures underwent decompaction, and minuscule protrusive structures were visible in almost all the spheroids (Fig. 3A). By day 21, morphological differences across the three constructs became rather pronounced. Under RGD+/TGF+ conditions, all spheroids were transformed into a dense, interconnected cellular network, with F-actin filaments stretched along the entire cellular structure (Fig. 3B). Notably, these cells displayed higher invasive capacity with strand-like single-cell migration (Fig. S9). Individually dispersed rounded cells or multicellular spheroids were not detected in these cultures. The RGD+/TGF- cultures yielded heterogeneous structures with varying sizes and morphologies, some small and compact, some appear to have loosened, while others displayed small, F-actin-rich filaments, invadopodium-like protrusions; however, extensive cell dissemination and invasion into the matrix was not seen. In the absence of RGD, and supplementation of TGFβ1 (RGD-/TGF+), cells formed compact spherical structures with cortical F-actin and no sign of invasion by day 21 of culture (Fig. 3B).

Inhibition studies were performed using SB-431542, a potent and selective inhibitor of TGFBR1/ALK5, and its relatives ALK4 and ALK7. It is known to inhibit TGFβ1 induced EMT and invasion in several cancers^30, 31^. The addition of SB-431542 (10 µM) substantially compromised the ability of cells to invade the matrix in RGD+/TGF+ cultures; cells now remained closely associated in the multicellular structures with protrusions. However, spheroids did not form the dense, interconnected cellular network in the presence of SB-431542 (Fig. 3C). On the other hand, SB-431542 did not significantly alter the morphology of the multicellular structures grown under RGD-/TGF+ and RGD+/TGF- conditions (Fig. 3C).

Cells residing in gels with or without RGD or TGFβ1 maintained high viability throughout the culture, as revealed by Live/dead assay (Fig. 4A). Morphological features observed under each condition were distinctly different, in agreement with our observations described in Fig. 3A, B. Quantitatively, cell viability was > 81% on day 1 and increased moderately to > 86% by day 21 for all three cultures (Fig. 4B, Fig. S10). PrestoBlue assay detected a significant (p < 0.05) increase in the number of metabolically active cells from day 1 to day 5 in RGD-/TGF+ cultures; no significant changes were observed thereafter. The RGD+/TGF- cultures displayed an initial increase in cell metabolism on day 10, followed by a significant (p < 0.05) decrease by day 21. A similar trend was observed in RGD+/TGF+ cultures, although cell metabolism peaked on day 5 (Fig. 4C).

**Fig. 4.**
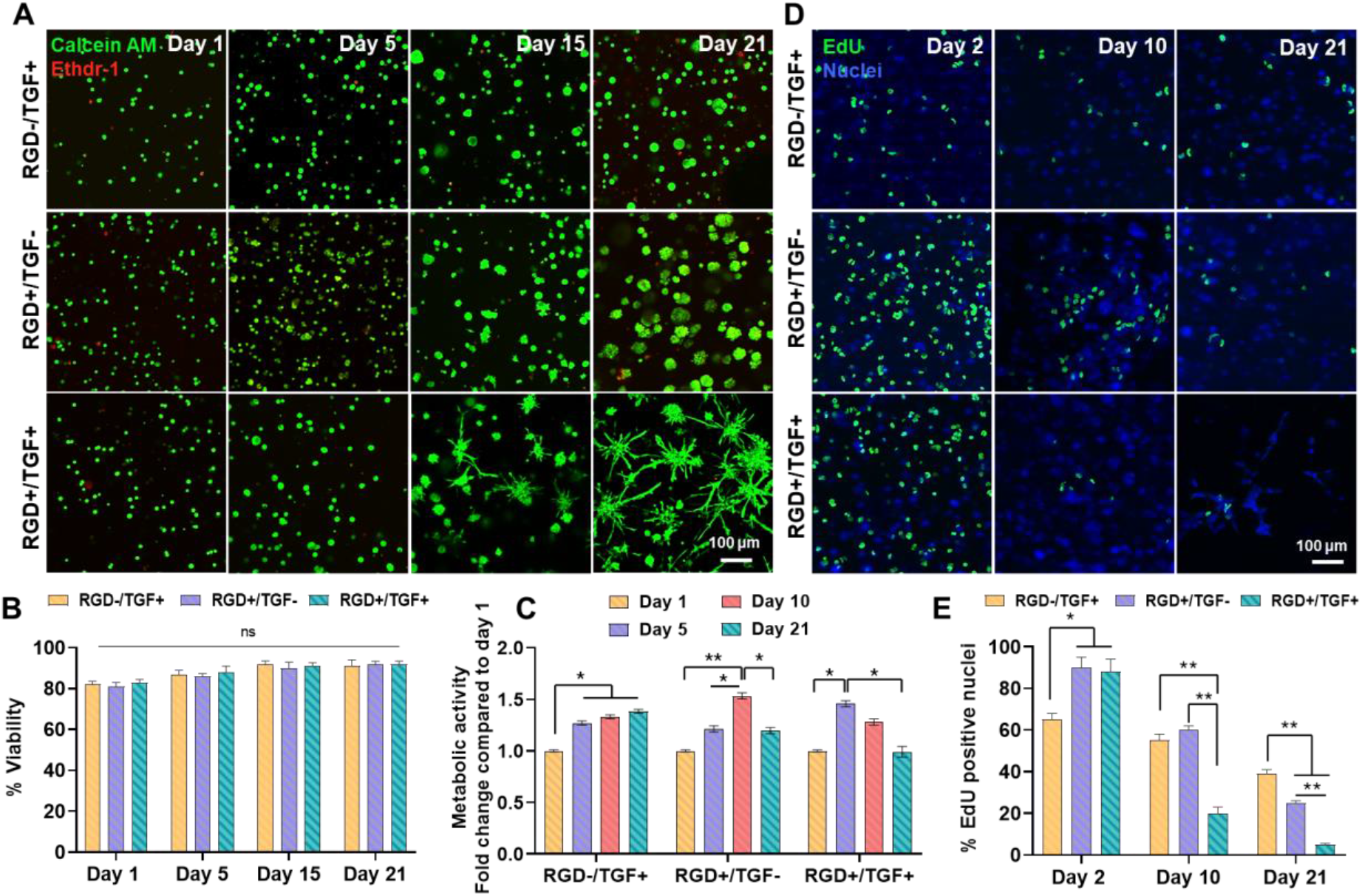
Characterization of 3D cultures by Live/dead, PrestoBlue and EdU assays. Cells were maintained in gels with 1.0 mM RGD in the presence of 10 ng/mL TGFβ1. Cultures with RGD or TGFβ1 only were included as the controls. **(A)** Representative confocal images of DU145 cells stained with calcein AM (green-live cells) and ethidium homodimer-1 (red-dead cells) after 1, 5, 15, and 21 days of culture. **(B)** Quantification of percent viability based on live/dead staining. Cell nuclei were counter-stained with Hoechst for quantification purposes. Error bars represent SEM. ns: not significant. **(C)** Quantification of cell viability/metabolic activity by PrestoBlue assay. The metabolic activity, as fold change, was normalized to day 1 of respective cultures. Error bars represent SEM. * p<0.05, ** p<0.01. **(D)** Detection of proliferating DU145 cells with Click-It EdU assay. EdU positive cells were detected with Alexa Fluor^TM^ 488 azide (green) and nuclei were counterstained with Hoechst (blue). Scale bar: 100 µm. **(E)** Quantification of percent EdU positive nuclei. Error bars represent SEM. * p<0.05, ** p<0.01.

Edu assay was performed to trace cell cycle kinetics by visualizing actively dividing cells (Fig. 4D, E). On day 2 of culture, that is, 24 h post-RGD conjugation, but before the addition of TGFβ1, the percent EdU positive nuclei was significantly (p < 0.05) higher (88-90%) in RGD cultures than the corresponding RGD-free controls (65%), confirming the pro-proliferative properties of immobilized RGD peptide. By day 10, percent EdU+ nuclei reduced significantly (p < 0.01) to 20 ± 3% in RGD+/TGF+ as compared to RGD-/TGF+ and RGD+/TGF- cultures. By day 21, percent EdU+ nuclei decreased across all three constructs; while over 40% of cells in RGD-/TGF+ were EdU positive, only 25 ± 1% in RGD+/TGF- and 5 ± 0.5 % cells in RGD+/TGF+ were EdU positive. Thus, the anti-proliferative effects of TGFβ1 were most profoundly manifested in RGD-containing cultures (Fig. 4E). A decrease in cell division could be indicative of cells adopting more aggressive and migratory phenotypes.

### RGD and TGFβ1 cooperatively activated TGFβ1 signaling pathways to induce EMT

We hypothesized that morphological changes seen in the TGFβ1-conditioned 3D cultures were associated with phenotypic alterations due to the activation of TGFβ signaling that is dependent on integrin-mediated adhesion to RGD. Using an RT2 profiler PCR array system, we first surveyed a panel of 84 known TGF target genes for day 21 cultures (Fig. 5A). Compared to the RGD+/TGF- controls, RGD+/TGF+ cultures exhibited a significant increase in the expression of *ENG*, *FN1*, *MAPK14*, *MMP2*, *SNAI1*, *TGFB2* and *TGFBR2*, all of which involved in cell differentiation and tissue development. A significant upregulation of a wide range of genes involved in cellular invasion and migration (*ACTA2*, *BDNF*, *IL10*, *KLF10*, *PDGFA*, *PTGS2*, *PTK2B*, *SERPINE1*, and *THBS1*) was also observed. In addition, the RGD+/TGF+ cultures exhibited elevated levels of cell cycle regulator *RHOB* and angiogenesis factor *VEGF*. Genes related to cell apoptosis that were upregulated included *GADD45B*, *MAP3K7*, and *MAPK8*. Genes involved in signal transduction pathways, including *ACVRL1*, *GLI2*, *HES1*, *ID1*, *ID2* and *ID3*, *RHOA*, *SMAD3*, *SMAD5* and *SMAD6*, also showed significant upregulation in these cultures. Genes encoding important transcription factors, including *AR* and *BHLHE40*, increased moderately. These results show that RGD conjugation is necessary to induce changes in TGF target genes. Alternatively, the array results from RGD+/TGF+ cultures were normalized to the RGD-/TGF+ counterparts. Interestingly, the gene expression profiles in both cases (i.e., normalization against RGD-/TGF+ vs normalization against RGD+/TGF-) were highly overlapping, indicating that integrin engagement via the immobilized RGD ligands primed the cancer cells to respond to the activation of the TGFβ-related pathways.

**Fig. 5.**
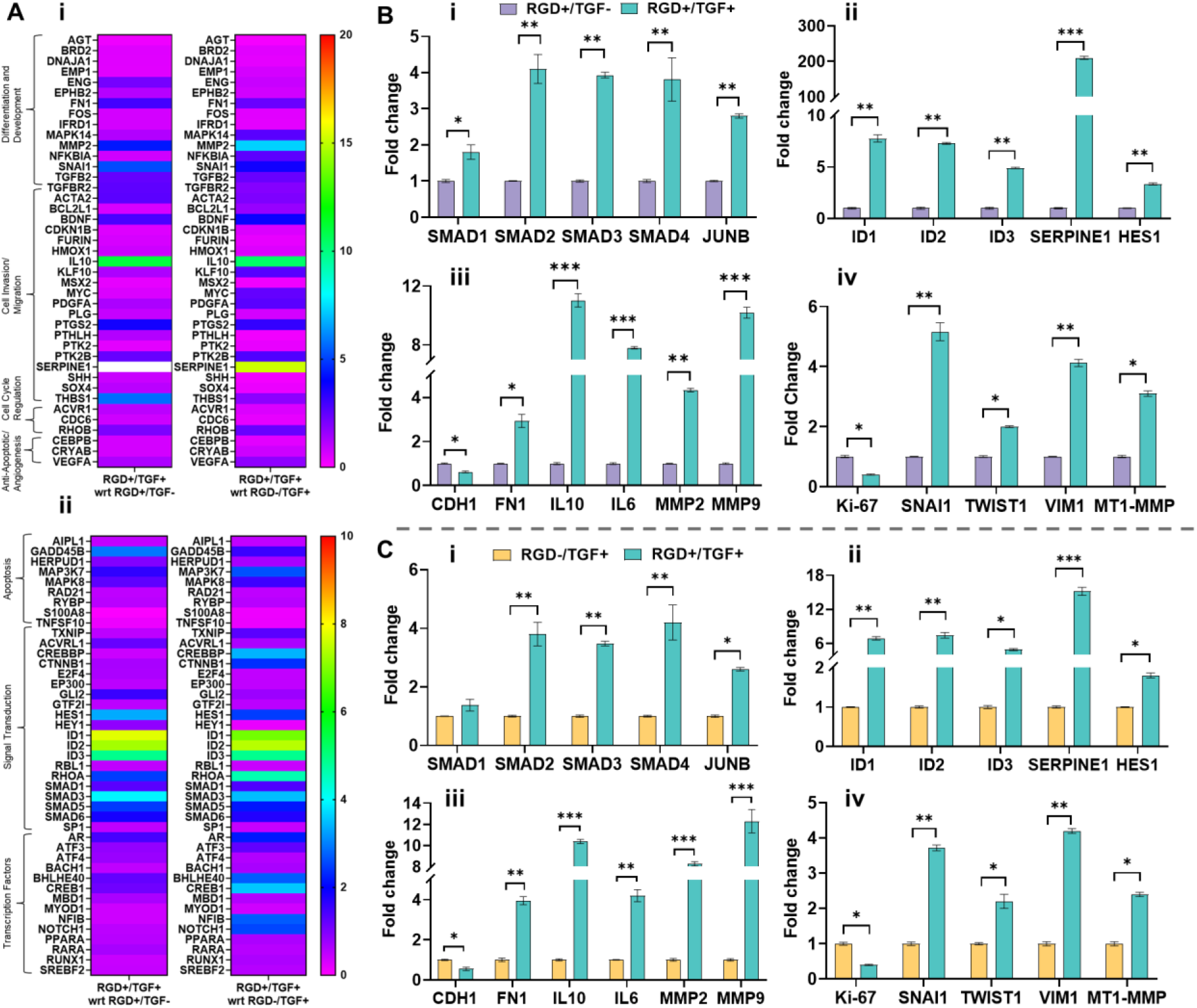
RT2 PCR profiler array (A) and RT qPCR (B-C) results detailing the expression of genes related to TGFβ1 targets, TGFβ1 signaling, and EMT. **(A)** Heat map depicting the transcriptional variations of all TGF target genes. The RT2 PCR array contains 84 TGF target genes (Tables S1-S7), 5 housekeeping genes including β-actin (*ACTB*), β-2-macroglobulin (*B2M*), glyceraldehyde 3-phosphate dehydrogenase (*GAPDH*), hypoxanthine phosphoribosyl transferase 1 (*HPRT1*), ribosomal protein lateral stalk subunit P0 (*RPLP0*), along with genomic DNA controls, reverse transcription controls and positive controls. Fold change was normalized TGFβ1 only (i) or RGD only (ii) cultures. **(B-C)** Fold change as analyzed by RT qPCR. Analysis was performed in RGD+/TGF+ constructs on day 21, and data was normalized to RGD+/TGF- (B) or RGD-/TGF+ (C). Error bars represent SEM, n=3, * p < 0.05, ** p < 0.01, *** p < 0.001. β-actin was used as a reference gene.

Cultures were further inspected quantitatively by RT qPCR (Fig. 5B-C). In agreement with the array results, TGFβ1-treated cultures showed a marked increase in classical TGFβ signaling pathway-related genes. Compared to the RGD+/TGF- controls, RGD+/TGF+ cultures (Fig. 5B) showed a significant upregulation of the expression of *SMAD1* (1.80 ± 0.20 fold), *SMAD2* (4.10 ± 0.4 fold), *SMAD3* (3.93 ± 0.08 fold), and *SMAD4* (3.81 ± 0.60 fold). A significant upregulation in the expression of the proto-oncogene *JUNB* (2.80 ± 0.06 fold) and tumor-promoting genes, including *ID1, ID2, ID3*, and *HES1*, along with over 200-fold increase in the expression of protease regulatory gene *SERPINE1* confirmed the stimulatory effects of TGFβ1. In the late stages of cancer progression, TGFβ1 functions as a tumor promoter by inducing EMT^32, 33^. Compared to the RGD+/TGF- controls, the RGD+/TGF+ cultures exhibited a significant decrease in the expression of *CDH1* (0.62 ± 0.04 fold) that encodes E-cadherin, an epithelial cell phenotypic marker, with a concomitant increase in the expression of *VIM1* (4.12 ± 0.12 fold) that encodes vimentin, a mesenchymal cell phenotype marker. EMT transcription factors *SNAI1* and *TWIST* were upregulated by 5.16 ± 0.30 and 2.00 ± 0.03 folds, respectively. Gene expressions of *MMP2, MMP9*, and *MT1-MMP*, the key players in ECM degradation, were increased in RGD+/TGF+ cultures by 4.33 ± 0.08, 10.20 ± 0.37, and 3.10 ± 0.09 folds, respectively. While the mRNA level of *FN1* was significantly (*p < 0.05*) increased (2.93 ± 0.30 fold), the expression of a cell proliferation marker *Ki-67* was significantly (*p < 0.05*) downregulated (0.40 ± 0.02 fold), in agreement with the EdU results. TGFβ1 treatment also significantly (*p < 0.001*) enhanced gene expressions of pro- inflammatory interleukins such as *IL10* (11.03 ± 0.45 fold) and *IL6* (7.80 ± 0.08 fold), consistent with previous observations^34^. A comparison was also made by normalizing the RGD+/TGF+ cultures against the RGD-/TGF+ counterparts (Fig. 5C). Again, the gene expression profile aligned well with that normalized against the RGD+/TGF- controls, although the specific fold changes varied from gene to gene. Collectively, our results confirmed that 3D culture of DU145 cells in RGD-conjugated, MMP-degradable HA gels in the presence of soluble TGFβ1 led to the upregulation of genes related to TGFβ signaling and EMT.

Next, cultures were examined by immunofluorescence to confirm the activation of SMAD- dependent TGFβ signaling and ascertain the acquisition of EMT at the protein level. In 2D cultures, nuclear localization of pSMAD3 was detected as early as 30 min after TGFβ1 stimulation, and thereafter, the signal intensity reduced, although it remained sequestered in the nucleus for up to 48 h. In the absence of TGFβ1, however, pSMAD3 was not detected (Fig. S11). In 3D, nuclear localization of pSMAD3 was detected on day three, 30 min after TGFβ1 was introduced (Fig. S12) in both TGFβ1-conditioned cultures (RGD-/TGF+ and RGD+/TGF+); however, pSMAD3 was not detected in cultures without TGFβ1 (i.e., RGD+/TGF-), confirming the inability of the 3D hydrogel alone to activate SMAD-dependent signaling. Although multicellular structures found in RGD-/TGF+ and RGD+/TGF+ cultures by day 21 exhibited disparate morphologies, both cultures showed positive staining for pSMAD3, albeit the signal was brighter in those with both RGD and TGFβ1 (Fig. 6A). Even with the immobilized RGD ligands, pSMAD3 signals were undetectable in RGD+/TGF- samples. SB-431542 supplementation abrogated pSMAD3 signals in RGD-/TGF+ and RGD+/TGF+ cultures (Fig. 6B), confirming the involvement of TGFβ1 in the activation of the SMAD-dependent signaling pathways. Similarly, JunB, an oncogenic transcription factor and TGFβ1 signaling target, was stained brightly in RGD+/TGF+ cultures but was not detected in RGD+/TGF- cultures. Noteworthy, in RGD-/TGF+ cultures, only 1-2 cells within the spheroids (white arrow, Fig. 6C) were stained positively for JunB. Collectively, these results indicated that TGFβ signaling was activated upon TGFβ1 binding to its receptor via the SMAD-dependent pathway. These results confirm that TGFβ1 is required to activate SMAD-dependent TGF signaling.

**Fig. 6.**
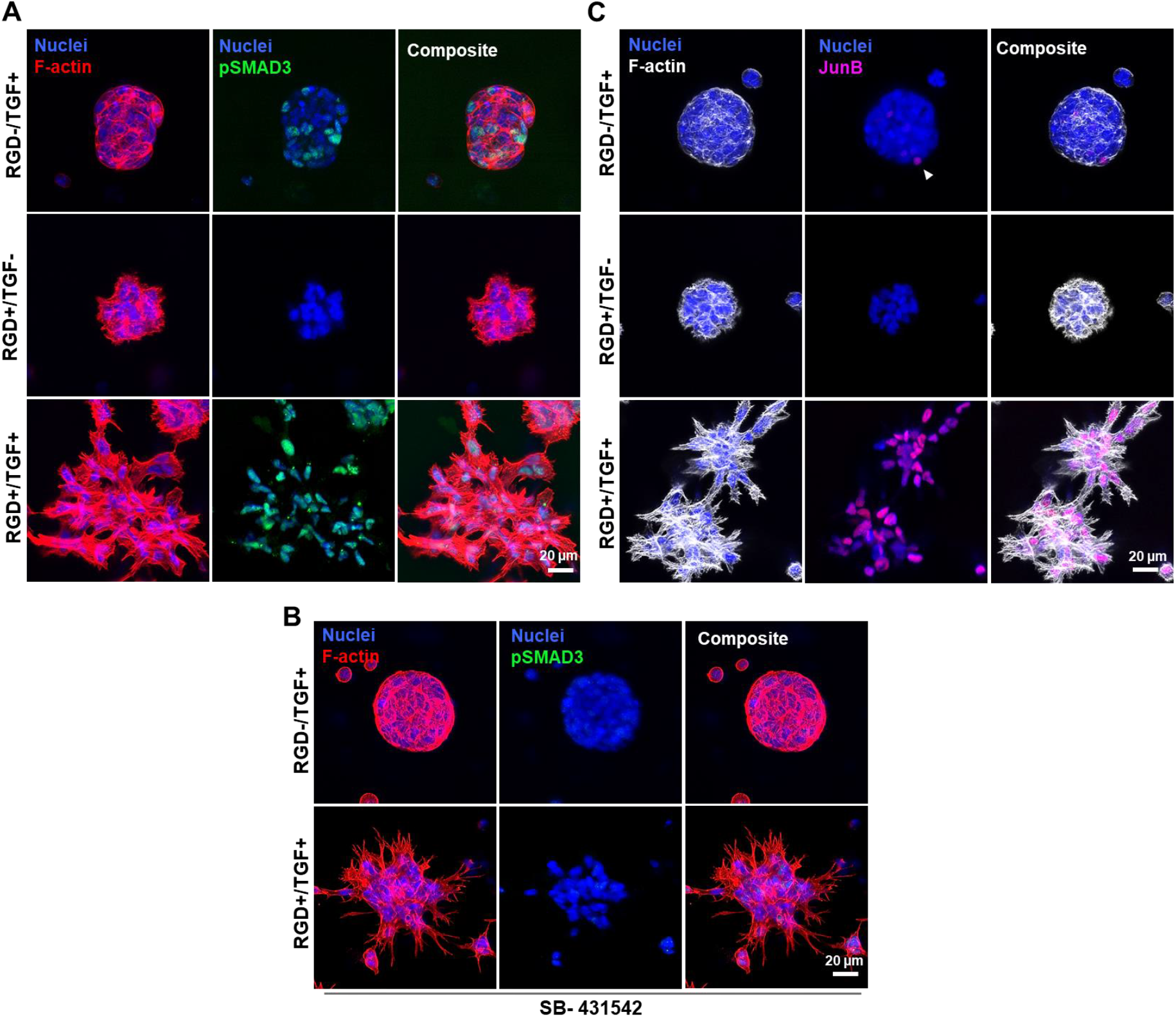
Representative confocal images of day 21 of 3D cultures confirming the activation of TGFβ1 signaling. **(A-B)** Cultures were maintained in the absence **(A)** or presence of SB-431542, a TGFβR1/ALK5 inhibitor **(B)**, and constructs were stained for pSMAD3 (green). Cell nuclei were counter-stained blue, and F-actin appeared red. **(C)** Cultures were maintained without SB-431542, and constructs were stained for JunB (magenta). Cell nuclei were counter-stained blue, and F-actin appeared gray. The white arrowhead indicates few nuclei stained for JunB. Scale bar: 20 µm.

Next, expression of EMT markers was assessed at the protein level by immunofluorescence for day 21 cultures. Under RGD-/TGF+ conditions, spheroids were stained brightly for adherens junction marker E-cadherin (Fig. 7A) and tight junctions marker ZO-1 (Fig. 7B) at the cell-cell junctions. The less compact spheroids developed under RGD+/TGF- conditions were stained weakly for E-cadherin and ZO-1, and the signals were rather diffuse and punctate. Contrarily, the highly elongated cells established in RGD+/TGF+ exhibited a complete depletion of cell-cell junction proteins, indicating a total loss of epithelial cell phenotype. As seen in (Fig. 7C), in RGD-/TGF+ cultures, only cells at the periphery of the spheroids interacting with the synthetic ECM were stained positive for vimentin, an intermediate type III filament present in mesenchymal cells. A similar staining pattern was observed under RGD+/TGF- conditions although the multicellular structures were less compact. In sharp contrast, the invading, dense cellular networks developed under RGD+/TGF+ conditions were intensely stained for vimentin. Finally, day 21 constructs were stained for classical epithelial cell markers, cytokeratins 18 (Krt18) and 19 (Krt19) (Fig. 8). The gain of vimentin expression corresponded to the loss of epithelial intermediate filaments. In both TGF-only and RGD-only cultures, cells in the spheroids with or without the invasive microstructures were stained brightly for both keratins. Contrarily, Krt18 and Krt19 signals were absent in RGD+/TGF+ cultures.. With SB-431542 treatment, Krt 18 and Krt19 expression was rescued in RGD+/TGF+ cultures. Collectively, TGFβ1 induced an aggressive mesenchymal phenotype via SMAD-dependent signaling pathways.

**Fig. 7.**
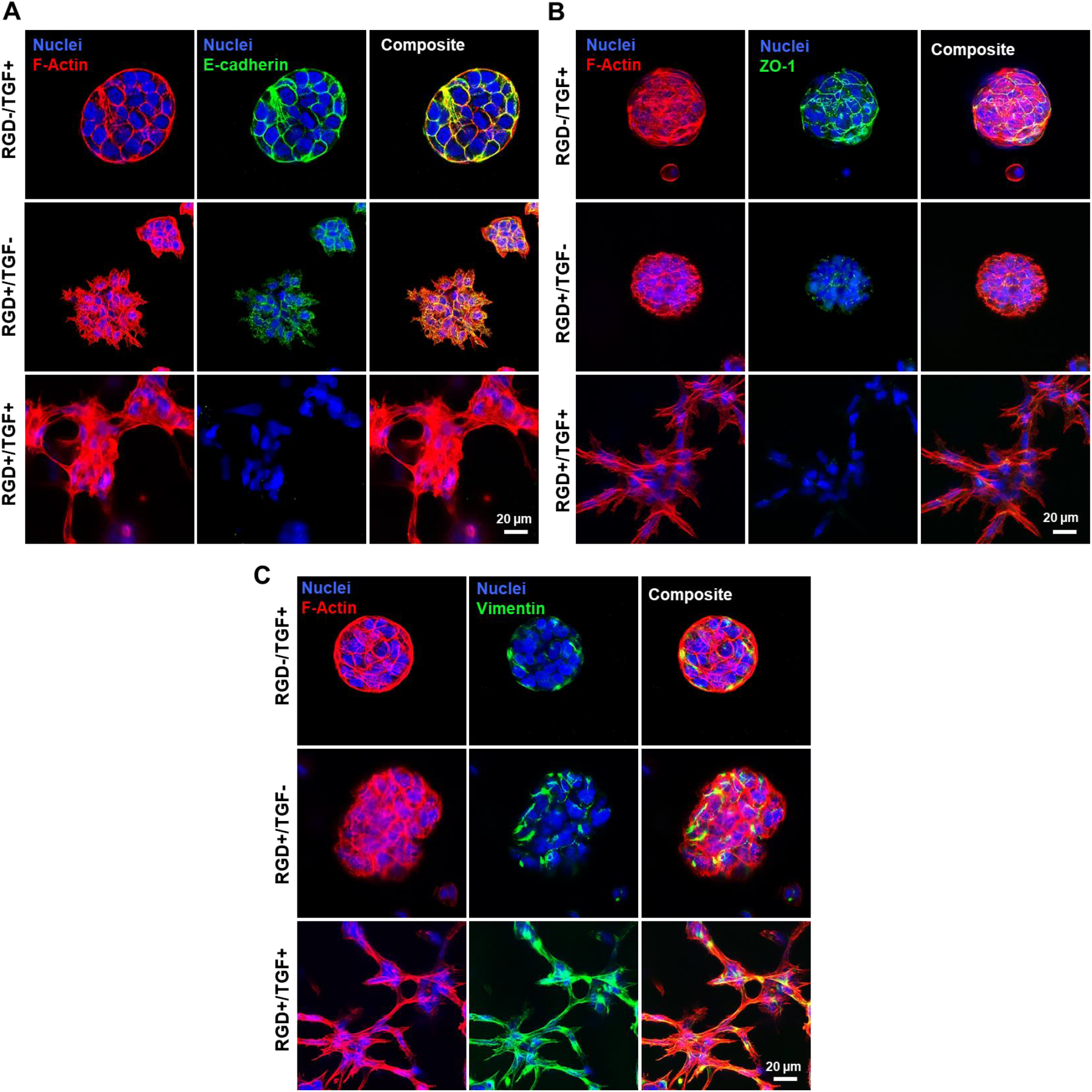
Representative immunofluorescent images confirming TGFβ1 induced EMT in DU145 cells. **(A)** Day 21 cultures were stained for the epithelial cell marker, E-cadherin (green). Nuclei and F-actin appeared blue and red, respectively. **(B)** Day 21 cultures were stained for the epithelial cell marker, ZO- 1 (green). Nuclei and F-actin appeared blue and red, respectively. **(C)** Day 21 cultures were stained for the mesenchymal cell marker, vimentin (green). Nuclei and F-actin appeared blue and red, respectively. Scale bar: 20 µm.

**Fig. 8.**
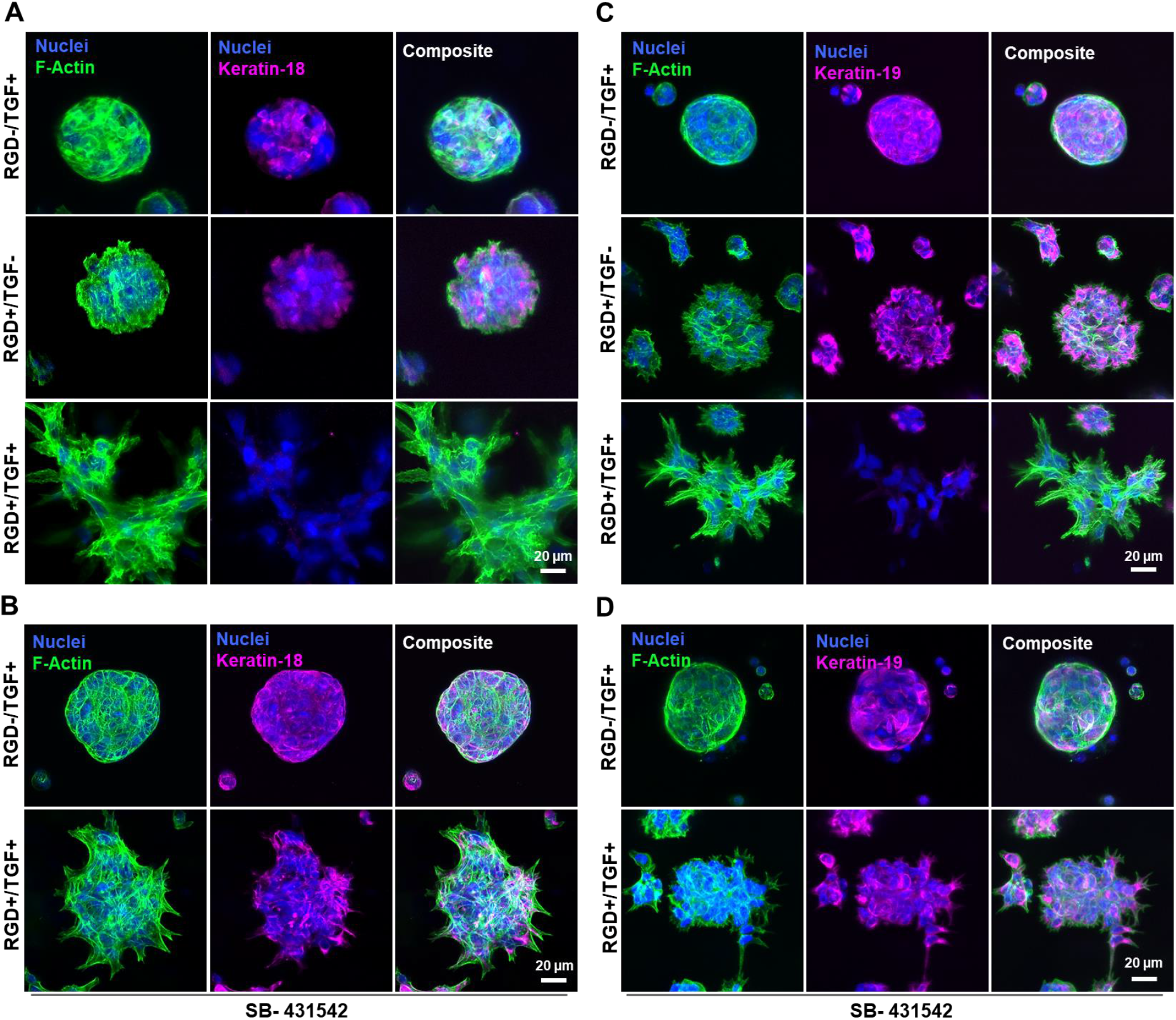
Representative confocal images of day 21 confirming TGFβ1 induced EMT in DU145 cells. **(A, B)** Cells were cultured in the absence (A) or presence (B) of SB-431542, a TGFβR1/ALK5 inhibitor, and the constructs were stained for epithelial cell marker Krt 18 (magenta). Cell nuclei were stained blue, and F-actin was stained green. **(C, D)** Cells were cultured in the absence (C) or presence (D) of SB-431542, and the constructs were stained for epithelial cell marker Krt 19 (magenta). Cell nuclei were stained blue and F-actin was stained green. Scale bar: 20 µm.

## DISCUSSION

Previously, we cultured DU145 cells in an MMP degradable HA-hydrogel that was rendered cell adhesive via bioorthogonal RGD tagging after compact, multicellular tumoroids had formed. In response to increased matrix adhesiveness, PCa cells underwent EMT by loosening cell-cell adhesion and strengthening cell-ECM interactions^24^. Herein, using the same hydrogel platform but with the cell adhesive ligands introduced earlier (day 1), we investigate how matrix properties, namely degradability and adhesiveness, direct cellular responses to a pro-oncogenic growth factor, TGFβ1. Cells maintained under all conditions exhibited high viability. Our short-term 2D scratch assay confirmed the ability of TGFβ1 to induce cell migration, potentially through the development of actin-rich protrusions at the leading edge and myosin motor-mediated retraction of the trailing edge^35, 36^

During cancer progression, the native ECM is largely remodeled by secreted MMPs, and excessive ECM degradation facilitates tumor invasion and metastasis^37–40^. When cultured in a nanoporous hydrogel resistant to MMP-mediated degradation, even in the presence of RGD and TGFβ1 signals, cells could not invade the matrix, although multicellular spheroids developed readily. In agreement with our prior observations^41–43^, the turnover of the HA hydrogel by cell-secreted hyaluronidase did not lead to morphological and phenotypic changes. Morphological changes in 3D are associated with local and global cell migration, facilitated by cell attachment to the matrix and MMP-mediated matrix degradation to create permissive ‘paths’^44–47^. Earlier work revealed invasion and metastasis of hepatocellular carcinoma were dependent on the reciprocal relationship between MMP-8 and TGFβ1^48^. Unlike the scrambled peptide crosslinker, SMR-bisNb used in this study is known to be degraded by MMP1, MMP2, MMP7, MMP9, and MT1-MMP^39^. When DU145 cells were cultured in MMP-degradable HA gels with a low RGD density (0.1, 0.3, and 0.5 mM), even in the presence of soluble TGFβ1, spheroids mostly remained compact, and wide-spread cell invasion into the matrix was not detected. Profound morphological changes were observed only when RGD concentration reached 1.0 mM; spheroids were transformed into elongated single cells with extensive cell-matrix interactions. Without TGFβ1, however, cells remained as loosened tumoroids with few protrusive structures. Without RGD, on the other hand, TGFβ1 did not induce any protrusion, and spheroids remained compact. Devoid of the adhesive sites, DU145 most likely behaved like epithelial cells, proliferating readily and maintaining energetically efficient round morphology with close cell-cell contacts^49–51^. Collectively, in the protease degradable matrix, the stimulatory effects of TGFβ1 were dependent on integrin binding with the RGD ligand. In other words, significant integrin engagement with the synthetic ECM is required to activate TGFβ1 signaling.

We examined the growth and differentiation of DU145 cells in MMP-degradable HA gels with or without RGD (1.0 mM) or TGFβ1 (10 ng/mL). It is well established that cell adhesion to matrices via integrin is indispensable for maintaining cells in a metabolically active state^43^. Moreover, TGFβ1 signaling can control cancer metabolic reprogramming and act as a metabolic driver to favor invasion and metastasis^52^. The PrestoBlue assay showed that RGD-containing cultures were metabolically more active on day 5 or 10 than on day 21. The EdU assay revealed a significantly higher percentage of proliferating cells on day 2 prior to the introduction of TGFβ1 and a significantly lower percentage of proliferating cells on day 21, confirming the anti-proliferative nature of TGFβ1^53^. The anti-proliferation effect of TGFβ1 is more pronouncedly manifested in RGD+ conditions. Therefore, TGFβ1 and RGD cooperatively stimulated morphological changes and phenotypic changes rather than increasing proliferation^54^. The EdU results align with the qPCR analysis for a proliferation marker Ki-67.

We showed that the regulatory functions of TGFβ1 were RGD-dependent. The upregulation of the expression of *TGFBR2* in RGD+/TGF+ cultures was likely due to TGFβ1 binding to its receptors TGFBR1/2. TGFBR2 plays a vital role in initiating downstream TGFβ1 signaling in tumorigenesis, and its expression is often altered in malignancies^55, 56^. Myeloid cells derived from patients with high-grade carcinoma showed increased expression of *TGFBR2*; genetic deletion of *TGFBR2* in myeloid cells significantly reduced cancer metastasis in mice^57^. The increased expression of *MAPK14* in RGD+/TGF+ cultures implies cells undergoing differentiation and engaging in ECM remodeling^58^. MAPK14 expression was associated with clinical stages of colorectal cancer, and patients with higher MAPK14 expression had poor prognosis and greater recurrence^59^.

RGD+/TGF+ cultures also exhibited upregulation of various genes involved in cell migration, invasion, and metastasis, including *ACTA2*, *BDNF*, *KLF10*, *PDGF*, *THBS1*, *VEGF,* and *RHOB*. Clinically, increased expression of ACTA2 is positively correlated to brain metastasis in lung adenocarcinoma^60^ and distant metastasis in EGFR-positive breast cancer^61^. BDNF is typically overexpressed in metastatic prostate, breast, and lung cancers^62^, while KLF10 is overexpressed in pancreatic cancer^63^. High expressions of PDGFA are associated with TGFβ1 induced EMT in breast cancer^64^, and high expressions of THBS1 lead to oral squamous cell carcinoma invasion by stimulating TGFβ1 induced expression of MMPs via integrin signaling^65^. Higher expression of VEGF, a proangiogenic factor, is correlated to increased metastatic potential in prostate cancer^66^. Additionally, the upregulation of RHOB, a Ras homolog gene family member involved in cell cycle regulation, was linked to the enhancement in cell motility and migration^67^. The upregulation of all these genes suggests that our 3D culture platform recapitulates the key features of clinically observed metastatic cancers.

Additionally, the expressions of key mediators of the canonical TGFβ1 signaling pathway, including *SMAD2*, *SMAD3*, and *SMAD4*^68^, were increased in RGD+/TGF+ cultures when compared to RGD-/TGF+ and RGD+/TGF- controls. Curiously, the expressions for SMAD5 and SMAD6 were also increased, although they do not play any major role in TGFβ1-induced cancer aggressiveness^69, 70^. TGFβ1 targets, including *JUNB, ID1, ID2, ID3, SERPINE1* and *HES1*, all showed higher expressions in RGD+/TGF+ cultures. As a member of the activator protein-1 (AP- 1) transcription factor, JunB is known to be overexpressed in many cancers and its knockout/knockdown inhibited cancer migration and invasion^71, 72^. Inhibitor of DNA binding (ID) proteins, a family of highly conserved transcriptional regulators^73^, play essential roles in triggering SMAD-dependent EMT in pre-malignant prostate epithelial progenitor cells^74^. Recent studies show that TGFβ1 treatment induced the expression of SERPINE1 in lung cancer cells, and numerous clinical studies reported the pro-tumorigenic effect of SERPINE1^75^. The prolonged presence of TGFβ1 from day 3 to day 21 did not lead to desensitization. Upon ligand binding, the TGFBRs remain active for several hours. However, repeated receptor activation maintains the entire SMAD2/3/4 complex within the nucleus for longer durations^76^. Protein-level characterizations by immunofluorescence further substantiated the gene-level observations. The marked increase in fluorescent signals detected in the nuclei for pSMAD3 and JunB in almost all cells in RGD+/TGF+ cultures is contrasted with faint or negative staining seen in TGFβ1 or RGD-only controls, respectively.

TGFβ1 promotes EMT by activating the SMAD-dependent pathways, leading to intense changes in cellular morphology, polarity, and phenotype^77^. In line with previous observations^78^, prolonged exposure of RGD+ 3D cultures to TGFβ1 induced EMT, as evidenced by substantial upregulation of type III intermediate filament and mesenchymal marker vimentin (*VIM1*) and downregulation of adherens junction protein E-cadherin (*CDH1*) and tight junction protein ZO-1, at both transcript and the protein levels. The upregulation of *SNAI1* and *TWIST* at the transcript levels further substantiates this claim. SNAI1 is a known transcriptional repressor of E-cadherin, where the loss of E-cadherin corresponds to increased SNAI1 expression. TWIST^79^, an indirect repressor of E-cadherin, can also promote metastasis and confer cancer cells with cancer stem cell-like properties, thereby increasing chemoresistance. The gain of mesenchymal cell markers like vimentin also correlated with the loss of other epithelial cell markers, such as cytokeratin 18/19, indicative of EMT^80^. Cytokeratin ensures the localization of E-cadherin to cell-cell junctions, and upon its disrupted expression, E-cadherin is mislocalized towards the basolateral membrane, and cells lose polarity^81^. In breast epithelial cells, Krt18 promotes nuclear localization and activation of SMAD2/3, and loss of Krt18 is accompanied by loss of epithelial markers and gain of mesenchymal markers^82^.

The RGD-/TGF+ and RGD+/TGF- cultures exhibited robust and consistent expression of epithelial cell markers Krt 18/19, although the staining was brighter for RGD-/TGF+ than for RGD+/TGF-. In both cases, peripheral cells interacting with the matrix were vimentin-positive, suggesting a hybrid EMT stage with mixed epithelial (interior) / mesenchymal (exterior) cell population within the spheroid. In RGD+/TGF+ cultures, complete loss of Krt 18/19 was accompanied by bright staining of mesenchymal marker, vimentin. Interestingly, in the absence of RGD, DU145 cells did not undergo EMT, even in the presence of TGFβ1. On the other hand, RGD alone is insufficient to confer widespread EMT with significant invasion into the matrix for almost all cells seen in the RGD+/TGF+ conditions. Our findings from the SB-431542 studies further corroborate our claim.

Collectively, our results established that TGFβ1-induced EMT requires integrin engagement with the synthetic ECM via RGD. Integrin binding can potentiate a TGF-dependent feed-forward loop that directs ECM remodeling, invasion, migration, and tumor progression. Because integrins are among the plethora of TGFβ regulated transcriptional targets, the interaction between TGFβ and integrins is bilateral^83^. Paradoxically, TGFβ1 can function as a tumor suppressor or promoter depending on the stages of cancer progression^84^. Our study suggests that changes in ECM composition can contribute to TGFβ1 role switching to function as a tumor promoter, thereby conferring widespread EMT.

## MATERIALS AND METHODS

### Maintenance of 2D Cell Culture

DU145 cells, a brain metastasized PCa cell line, were purchased from the American Type Culture Collection (ATCC, Manassas, VA). Green fluorescent protein (GFP)-labeled DU145 cells were acquired from Kenneth Pienta’s lab at Johns Hopkins Medical Institute. Cells were expanded in Eagle’s Minimum Essential Medium (EMEM) that was supplemented with 10% (v/v) fetal bovine serum (FBS, Cytiva HyClone, Logan, UT) and 1% (v/v) penicillin-streptomycin. Media was refreshed every 2 days and cultures were maintained at 37°C with 85% humidity and 5% CO_2_ for around 7 days until they reached 75-80% confluency. Cells were then treated with 0.25% (w/v) trypsin containing ethylenediaminetetraacetic acid (2.2 mM EDTA-4Na) for 3 min and residual trypsin was neutralized with an equal volume of EMEM. Cells were centrifuged at 150 g for 5 min, the supernatant was removed and resuspended in fresh EMEM to continue further passages. All experiments were conducted using passages 6-12.

### Scratch Assay

A two-well silicone insert (Ibidi USA Inc, Fitchburg, WI) with a defined cell-free gap of 500 µm was placed inside a Nunc™ Lab-Tek™ 2 Chamber Slide™ (Thermo Fisher Scientific, Waltham, MA). Seventy microliter EMEM containing 0.6x10^6^ GPF-tagged DU145 per mL was added to each insert well. Cells were allowed to attach for 24 h, after which the silicone insert was carefully removed to create a cell-free gap. Cells were replenished with EMEM supplemented with or without TGFβ1 at 10 ng/mL. Cell migration/wound closure was analyzed using Zeiss LSM 880 (Carl Zeiss, Oberkochen, Germany) by capturing images from the same location at 0, 12, 24, and 48 h with a C-Apochromat 5× air objective with 5 µm z-stacks with 1 µm z-axis step size. The brightness for each channel was uniformly adjusted using ZEN 3.0 SR software. Representative images were processed as maximum-intensity projections. Experiments were carried out using three biological repeats for each condition.

### Cell Encapsulation and 3D Culture

HA-Tz, RGD/RGE-TCO, and SMR-bisNb were synthesized and characterized following our previously reported methods^24–26^. SMV-bisNb (SMVGMRPG) ^28^ was prepared by adapting the procedure for SMR-bisNb. All hydrogel components were sterile filtered (with 0.22 µm filters) before use. To establish the cellular construct, HA-Tz, and SMR-bisNb or SMV-bisNb were dissolved separately in PBS at a stock concentration of 2 wt% and 10 mM, respectively. RGD-TCO and RGE-TCO were dissolved in EMEM at 4 mM. HA-Tz and SMR-bisNb were mixed at a Tz/Nb molar ratio of 2/1 to yield a mixture with the final Tz and Nb concentrations of 5.0 mM and 2.5 mM, respectively. DU145 cells were suspended in the above solution at 0.5 x 10^6^ cells/mL and 300 µL or 50 µL of the viscous solution was transferred to a 12 mm PTFE cell culture insert (Millicell 0.4 μm, EMD Millipore Corporation, Burlington, MA) or a 48-well glass-bottom MatTek plate (MatTek Corporation, Ashland, MA). The mixture was maintained at 37°C for 1 h to allow complete gelation before EMEM was added (Day 0). One day later (day 1), media containing 1.6 mM RGD-TCO and 2.4 mM RGE-TCO was added to the cell-laden hydrogel to initiate diffusion-controlled interfacial ligation at 37°C. Twenty-four hours later, the TCO reservoir was replaced with fresh EMEM. On day 3, TGFβ1 (PeproTech, Rocky Hill, NJ) was introduced at 10 ng/ml and cultures were maintained until day 21 in TGFβ1-conditioned media, with media refreshment every 3 days. Cultures maintained in gels without RGD or TGFβ1 were included as the controls. For TGFβRI inhibition experiments, SB-431542 (Tocris Bioscience, Bristol, UK,) was added at 10 μM to TGFβ1 conditioned media on day 3, and cultures were maintained with the inhibitor until day 21. Bright-field images were captured using an Eclipse Ti-E microscope (Nikon, Tokyo, Japan) on days 0,10 and 21.

### Live/dead Assay

Cell viability was analyzed using LIVE/DEAD^TM^ viability assay kit (Thermo Fisher Scientific). Calcein AM (1:1000 dilution), ethidium homodimer (EthD-1, 1:2000 dilution), and Hoechst 33342 (1:200 dilution) were diluted in warm 1X PBS. Constructs were incubated in the dye solution for 20 min at 37°C. After thorough washes with PBS (3 times), constructs were imaged using an LSM 880 confocal microscope with C-Apochromat 10×/0.45 NA water immersion objective. Images were captured from random sections of hydrogel as 150 µm z-stacks with z z-axis step size of 7 µm and processed as maximum intensity projections. A total of three biological replicates for each day point were used for capturing images, and at least ten images were used to calculate cell viability. Cell viability was calculated using ImageJ (National Institutes of Health, Bethesda, MD) as previously described.

### Presto Blue Assay

The metabolic activity of encapsulated DU145 cells in hydrogel constructs was assessed using the PrestoBlue reagent (Thermo Fisher Scientific). The reagent was diluted in warm EMEM at 10% (v/v). At a predetermined time, the media was replaced with the PrestoBlue solution, and the plate was covered with aluminum foil and maintained at 37°C for 3 h. Then, 100 µL solution from each hydrogel construct was transferred to a fresh 96-well plate, with 3 technical and 3 biological repeats per construct. Media blank was also included for each time point. Fluorescence was measured at 585 nm using Spectramax i3x Multimode Microplate Reader (Molecular Devices, San Jose, CA) with gentle plates shaking between each reading. Media blanks were subtracted from each reading, and fold change in metabolic activity was calculated by normalizing data to the day 1 reading for each condition.

### EdU Cell Proliferation Assay

Click-iT™ EdU Imaging Kit with Alexa Fluor™ 488 (Thermo Fisher Scientific) was used to characterize cell proliferation. A fresh EdU solution in warm EMEM was added to the media at 10 µM, and the construct was incubated at 37°C, with 85% humidity and 5% CO_2_ for 3 h. After the media was removed, the construct was fixed with 4% (w/v) paraformaldehyde (PFA, Sigma Aldrich, Burlington, MA) in PBS for 30 min. After two washes with 3% (w/v) bovine serum albumin (BSA) in PBS, the construct was incubated in 0.5% (v/v) Triton X-100 in PBS for 20 min to permeabilize the cells, followed by two washes with 3% BSA. The Click-iT^TM^ reaction cocktail was freshly prepared by mixing the Click-iT^TM^ reaction buffer, CuSO_4_, and Alexa Fluor®-488, following the manufacturer’s protocol. Upon removal of the BSA solution, the construct was incubated with 500 µL of Click-iT^TM^ reaction cocktail for 1 h with the cell culture plate covered with aluminum foil. The reaction cocktail was removed, and the construct was washed with 3% BSA two times. Finally, Hoechst 33342 (1:200) was used to counterstain the nuclei. Images were captured using an LSM 880 confocal microscope with C-Apochromat 10x/0.45 NA water immersion objective and airy scan detector with 100 µm z-stacks with 7 µm z-axis step size, each 16-bit. EdU-positive nuclei were counted using ImageJ using 10 images for each condition.

### RT2-Profiler PCR Array

Hydrogel constructs were thoroughly washed with PBS three times, snap-frozen on a dry ice/isopropanol mixture, and stored at -80°C till further use. Frozen samples were crushed with a pestle, and the gel slurry was dissociated using Trizol (Invitrogen, Carlsbad, CA). RNA extraction was performed following our reported method^25^, and the dry RNA pellet was reconstituted in 20 µL nuclease-free DEPC-treated water. RNA quantification and purity were assessed using a NanoDrop 2000 spectrophotometer (Nanodrop Technologies, Wilmington, DE). Using RT2 First Strand Kit (Qiagen, Hilden, Germany), high-quality cDNA was synthesized by reverse transcribing 0.5 µg RNA. The RT2 Profiler PCR Array (Qiagen, Hilden, Germany) in a format-A, 96-well plate was used to identify 84 differentially expressed genes related to TGF signaling and their targets. The PCR array was conducted using RT2 SYBR Green ROX qPCR Master mix (Qiagen), following the manufacturer’s protocol. Each of the 96-well plates consisted of 84 TGF target genes (Tables S1-S7), 5 housekeeping genes including β-actin (*ACTB*), β-2-macroglobulin (*B2M*), glyceraldehyde 3-phosphate dehydrogenase (*GAPDH*), hypoxanthine phosphoribosyl transferase 1 (*HPRT1*), ribosomal protein lateral stalk subunit P0 (*RPLP0*), along with genomic DNA controls, reverse transcription controls and positive controls. The average of these five housekeeping genes was used for calculating fold change using 2^-ΔΔCT^ method. The PCR array analysis was conducted normalized TGFβ1 only or RGD only cultures, and a heat map was generated to plot 84 TGF target genes. The heat map depicts the transcriptional variations of all TGF target genes belonging to different functional categories namely differentiation/development, cell invasion/migration, cell cycle regulation, anti-apoptosis/angiogenesis, apoptosis, signal transduction, and transcription factors.

### Real-Time Quantitative Polymerase Chain Reaction (RT-qPCR)

High-quality RNA was extracted following procedures described above. RNA samples with absorbance at λ 260/280 ratio of 1.9-2.1 and λ 260/230 ratio of 2.0-2.2 were used for reverse transcription. Using QuantiTect Reverse Transcription Kit (Qiagen, Hilden, Germany), 0.5 µg of RNA was reverse transcribed to cDNA following the manufacturer’s protocol. After combining Power SYBR green master mix (1/2, v/v), forward and reverse primers (Integrated DNA Technologies, Coralville, IA), cDNA template, and water, qPCR was performed using ABI 7300 real-time PCR system. β-actin was used as an endogenous reference gene, and fold change was calculated using the 2^-ΔΔCT^ method. A total of three biological repeats for each experimental condition were analyzed from three technical replicates. The forward and reverse primer pair information can be found in Table S8.

### Immunocytochemistry

After the cell culture media was aspirated, the construct was washed with 1X PBS three times and fixed with 4% (w/v) PFA for 30 min. To stain for pSMAD3 and JunB, constructs were permeabilized in 0.2% (v/v) Triton X-100 for 3 h and blocked in 3% (w/v) BSA for 3 h at room temperature. To stain for E-cadherin and ZO-1, samples were permeabilized in 0.1% Triton X-100 for 10 min, and blocked in 3% BSA for 16 h at 4°C. To stain for keratin 18,19, and vimentin, samples were permeabilized with 0.2% (v/v) Triton X-100 for 2 h at room temperature and blocked in 3% BSA for 16 h at 4°C. The permeabilized samples were incubated in primary antibody solutions with appropriate dilutions in 3% BSA overnight at 4°C for pSMAD3, JunB, E-cadherin, and ZO-1, for 6 h at room temperature for keratin 18,19 and for 2 h at room temperature for vimentin. Primary antibody details can be found in Table S9. After aspirating the primary antibody solution, constructs were washed with PBS twice, then twice with PBS containing 0.05% tween 20 (PBST, Sigma Aldrich), and again two times with PBS, with 15 min incubation for every washing step. Alexa Fluor 488 or 647 AffiniPure Fab Fragment Goat Anti-mouse or anti-rabbit IgG (H+L) secondary antibodies (Jackson Immunoresearch Labs, West Grove, PA) and Alexa Fluor^TM^ 568 incubated in the secondary antibody solution for 4 h at room temperature. The solution was aspirated, and the samples were washed with PBST and PBS three times each, with 15 min incubation at each step. Nuclei were counter-stained by 4′,6-diamidino-2- phenylindole (DAPI, Life Technologies), diluted at 1/1000 in PBS, for 45 min at room temperature. Finally, immunostained hydrogel constructs were washed with PBS for 24 h at room temperature. Antibody incubation and washing steps were conducted using an orbit shaker at 100 and 250 rpm, respectively. To stain for F-actin and nuclei only, hydrogel constructs were permeabilized in 0.2% (v/v) Triton X-100 for 2 h at room temperature and then stained with Alexa Fluor^TM^ 568 phalloidin and DAPI, as described above. Fluorescent microscopy was performed using Zeiss laser scanning microscope 880 (LSM880) using AiryScan detector in Fast Airy scan mode with either C-Apochromat 10×/0.45 water or LD LCI Plan-Apochromat 25×/0.8 Imm Korr water/oil objectives. Images were captured as 75 µm z-stacks with 1 µm z-axis step size, each 16-bit. After airy scan processing, images were saved as maximum intensity projection high-resolution images. The brightness for each channel was uniformly adjusted using ZEN 3.0 SR software for individual representative images. At least 3 biological replicates for each condition were used to capture approximately 15 images for each construct.

### Statistical Analysis

Statistical analysis was performed using one-way ANOVA followed by Tukey’s HSD post hoc for pairwise comparison or student’s t-test. A p-value of <0.05 was considered significantly different. Statistical analysis was conducted using a JMP Pro 15 (SAS Institute Inc) and plotted using GraphPad Prism (San Diego, CA) version 9.2.0. Error bars represent the standard error of the mean value (SEM). For all experiments, replicates were collected across different passages on different days, with at least three biological repeats. The sample size was selected based on preliminary experimental data to ensure sufficient statistical power.

## Acknowledgments Funding

National Institutes of Health grant R01 DE029655 (XJ, JMF) National Institutes of Health grant NIDCD, R01DC014461 (XJ, JMF) National Science Foundation DMR 1809612 (XJ, JMF)

National Science Foundation DMR 2011824 (XJ, JMF)

## Author contributions

Conceptualization: MP, XJ Methodology: MP, HG, JMF, XJ Investigation: MP, HG Visualization: MP

Supervision: XJ, JMF Writing—original draft: MP, HG

Writing—review & editing: MP, HG, JMF, XJ

## Data and materials availability

All data are available in the main text or the supplementary materials.

## REFERENCES

(1) Sung, H.; Ferlay, J.; Siegel, R. L.; Laversanne, M.; Soerjomataram, I.; Jemal, A.; Bray, F. Global Cancer Statistics 2020: GLOBOCAN Estimates of Incidence and Mortality Worldwide for 36 Cancers in 185 Countries. CA Cancer J Clin 2021, 71 (3), 209–249. DOI: 10.3322/caac.21660.

(2) Rebello, R. J.; Oing, C.; Knudsen, K. E.; Loeb, S.; Johnson, D. C.; Reiter, R. E.; Gillessen, S.; Van der Kwast, T.; Bristow, R. G. Prostate cancer. *Nat Rev Dis Primers* 2021, 7 (1), 9. DOI: 10.1038/s41572-020-00243-0.

(3) Siegel, R. L.; Miller, K. D.; Jemal, A. Cancer statistics, 2019. CA Cancer J Clin 2019, 69 (1), 7–34. DOI: 10.3322/caac.21551.

(4) Kumar, A.; Coleman, I.; Morrissey, C.; Zhang, X.; True, L. D.; Gulati, R.; Etzioni, R.; Bolouri, H.; Montgomery, B.; White, T.;, et al. Substantial interindividual and limited intraindividual genomic diversity among tumors from men with metastatic prostate cancer. Nat Med 2016, 22 (4), 369–378. DOI: 10.1038/nm.4053.

(5) Ali, A.; Du Feu, A.; Oliveira, P.; Choudhury, A.; Bristow, R. G.; Baena, E. Prostate zones and cancer: lost in transition? Nat Rev Urol 2022, 19 (2), 101–115. DOI: 10.1038/s41585-021-00524-7.

(6) Paget, S. The distribution of secondary growths in cancer of the breast. 1889. Cancer Metastasis Rev 1989, 8 (2), 98–101.

(7) Fidler, I. J. The pathogenesis of cancer metastasis: the ’seed and soil’ hypothesis revisited. Nat Rev Cancer 2003, 3 (6), 453–458. DOI: 10.1038/nrc1098.

(8) David, C. J.; Massagué, J. Contextual determinants of TGFβ action in development, immunity and cancer. Nat Rev Mol Cell Biol 2018, 19 (7), 419–435. DOI: 10.1038/s41580-018-0007-0.

(9) Shi, M.; Zhu, J.; Wang, R.; Chen, X.; Mi, L.; Walz, T.; Springer, T. A. Latent TGF-β structure and activation. Nature 2011, 474 (7351), 343–349. DOI: 10.1038/nature10152.

(10) Wu, M. Y.; Hill, C. S. Tgf-beta superfamily signaling in embryonic development and homeostasis. Dev Cell 2009, 16 (3), 329–343. DOI: 10.1016/j.devcel.2009.02.012.

(11) Taipale, J.; Saharinen, J.; Keski-Oja, J. Extracellular matrix-associated transforming growth factor-beta: role in cancer cell growth and invasion. Adv Cancer Res 1998, 75, 87–134. DOI: 10.1016/s0065-230x(08)60740-x.

(12) Derynck, R.; Zhang, Y. E. Smad-dependent and Smad-independent pathways in TGF-beta family signalling. Nature 2003, 425 (6958), 577–584. DOI: 10.1038/nature02006.

(13) Massagué, J. TGFbeta in Cancer. Cell 2008, 134 (2), 215–230. DOI: 10.1016/j.cell.2008.07.001.

(14) Siegel, P. M.; Massagué, J. Cytostatic and apoptotic actions of TGF-beta in homeostasis and cancer. Nat Rev Cancer 2003, 3 (11), 807–821. DOI: 10.1038/nrc1208.

(15) Massagué, J. G1 cell-cycle control and cancer. Nature 2004, 432 (7015), 298–306. DOI: 10.1038/nature03094.

(16) Miettinen, P. J.; Ebner, R.; Lopez, A. R.; Derynck, R. TGF-beta induced transdifferentiation of mammary epithelial cells to mesenchymal cells: involvement of type I receptors. J Cell Biol 1994, 127 (6 Pt 2), 2021-2036. DOI: 10.1083/jcb.127.6.2021.

(17) Caulín, C.; Scholl, F. G.; Frontelo, P.; Gamallo, C.; Quintanilla, M. Chronic exposure of cultured transformed mouse epidermal cells to transforming growth factor-beta 1 induces an epithelial-mesenchymal transdifferentiation and a spindle tumoral phenotype. Cell Growth Differ 1995, 6 (8), 1027–1035.

(18) Pastushenko, I.; Blanpain, C. EMT Transition States during Tumor Progression and Metastasis. Trends Cell Biol 2019, 29 (3), 212–226. DOI: 10.1016/j.tcb.2018.12.001.

(19) Arriaga, J. M.; Abate-Shen, C. Genetically Engineered Mouse Models of Prostate Cancer in the Postgenomic Era. Cold Spring Harb Perspect Med 2019, 9 (2). DOI: 10.1101/cshperspect.a030528.

(20) Risbridger, G. P.; Toivanen, R.; Taylor, R. A. Preclinical Models of Prostate Cancer: Patient-Derived Xenografts, Organoids, and Other Explant Models. Cold Spring Harb Perspect Med 2018, 8 (8). DOI: 10.1101/cshperspect.a030536.

(21) Ellem, S. J.; De-Juan-Pardo, E. M.; Risbridger, G. P. In vitro modeling of the prostate cancer microenvironment. Adv Drug Deliv Rev 2014, 79-80, 214–221. DOI: 10.1016/j.addr.2014.04.008.

(22) Baker, B. M.; Chen, C. S. Deconstructing the third dimension: how 3D culture microenvironments alter cellular cues. J Cell Sci 2012, 125 (Pt 13), 3015–3024. DOI: 10.1242/jcs.079509.

(23) Xu, X.; Farach-Carson, M. C.; Jia, X. Three-dimensional in vitro tumor models for cancer research and drug evaluation. Biotechnol Adv 2014, 32 (7), 1256–1268. DOI: 10.1016/j.biotechadv.2014.07.009.

(24) Pol, M.; Gao, H.; Zhang, H.; George, O. J.; Fox, J. M.; Jia, X. Dynamic modulation of matrix adhesiveness induces epithelial-to-mesenchymal transition in prostate cancer cells in 3D. Biomaterials 2023, 299, 122180. DOI: 10.1016/j.biomaterials.2023.122180.

(25) Gao, H.; Pol, M.; Makara, C. A.; Song, J.; Zhang, H.; Zou, X.; Benson, J. M.; Burris, D. L.; Fox, J. M.; Jia, X. Bio-orthogonal tuning of matrix properties during 3D cell culture to induce morphological and phenotypic changes. Nat Protoc 2024. DOI: 10.1038/s41596-024-01066-z.

(26) Song, J.; Gao, H.; Zhang, H.; George, O. J.; Hillman, A. S.; Fox, J. M.; Jia, X. Matrix Adhesiveness Regulates Myofibroblast Differentiation from Vocal Fold Fibroblasts in a Bio- orthogonally Cross-linked Hydrogel. ACS Appl Mater Interfaces 2022, 14 (46), 51669–51682. DOI: 10.1021/acsami.2c13852.

(27) Zou, X.; Zhang, H.; Benson, J. M.; Gao, H.; Burris, D. L.; Fox, J. M.; Jia, X. Modeling the Maturation of the Vocal Fold Lamina Propria Using a Bioorthogonally Tunable Hydrogel Platform. Adv Healthc Mater 2023, 12 (29), e2301701. DOI: 10.1002/adhm.202301701.

(28) Guenther, C. M.; Brun, M. J.; Bennett, A. D.; Ho, M. L.; Chen, W.; Zhu, B.; Lam, M.; Yamagami, M.; Kwon, S.; Bhattacharya, N.;, et al. Protease-Activatable Adeno-Associated Virus Vector for Gene Delivery to Damaged Heart Tissue. Mol Ther 2019, 27 (3), 611–622. DOI: 10.1016/j.ymthe.2019.01.015.

(29) Liu, Z.; Wang, L.; Xu, H.; Du, Q.; Li, L.; Zhang, E. S.; Chen, G.; Wang, Y. Heterogeneous Responses to Mechanical Force of Prostate Cancer Cells Inducing Different Metastasis Patterns. Adv Sci (Weinh) 2020, 7 (15), 1903583. DOI: 10.1002/advs.201903583.

(30) Halder, S. K.; Beauchamp, R. D.; Datta, P. K. A specific inhibitor of TGF-beta receptor kinase, SB-431542, as a potent antitumor agent for human cancers. Neoplasia 2005, 7 (5), 509–521. DOI: 10.1593/neo.04640.

(31) Inman, G. J.; Nicolás, F. J.; Callahan, J. F.; Harling, J. D.; Gaster, L. M.; Reith, A. D.; Laping, N. J.; Hill, C. S. SB-431542 is a potent and specific inhibitor of transforming growth factor-beta superfamily type I activin receptor-like kinase (ALK) receptors ALK4, ALK5, and ALK7. Mol Pharmacol 2002, 62 (1), 65–74. DOI: 10.1124/mol.62.1.65.

(32) Peinado, H.; Quintanilla, M.; Cano, A. Transforming growth factor beta-1 induces snail transcription factor in epithelial cell lines: mechanisms for epithelial mesenchymal transitions. J Biol Chem 2003, 278 (23), 21113–21123. DOI: 10.1074/jbc.M211304200.

(33) Hao, Y.; Baker, D.; Ten Dijke, P. TGF-β-Mediated Epithelial-Mesenchymal Transition and Cancer Metastasis. Int J Mol Sci 2019, 20 (11). DOI: 10.3390/ijms20112767.

(34) Park, J. I.; Lee, M. G.; Cho, K.; Park, B. J.; Chae, K. S.; Byun, D. S.; Ryu, B. K.; Park, Y. K.; Chi, S. G. Transforming growth factor-beta1 activates interleukin-6 expression in prostate cancer cells through the synergistic collaboration of the Smad2, p38-NF-kappaB, JNK, and Ras signaling pathways. Oncogene 2003, 22 (28), 4314–4332. DOI: 10.1038/sj.onc.1206478.

(35) Chi, Q.; Yin, T.; Gregersen, H.; Deng, X.; Fan, Y.; Zhao, J.; Liao, D.; Wang, G. Rear actomyosin contractility-driven directional cell migration in three-dimensional matrices: a mechano-chemical coupling mechanism. J R Soc Interface 2014, 11 (95), 20131072. DOI: 10.1098/rsif.2013.1072.

(36) Ridley, A. J.; Schwartz, M. A.; Burridge, K.; Firtel, R. A.; Ginsberg, M. H.; Borisy, G.; Parsons, J. T.; Horwitz, A. R. Cell migration: integrating signals from front to back. Science 2003, 302 (5651), 1704–1709. DOI: 10.1126/science.1092053.

(37) Saarialho-Kere, U. K.; Chang, E. S.; Welgus, H. G.; Parks, W. C. Distinct localization of collagenase and tissue inhibitor of metalloproteinases expression in wound healing associated with ulcerative pyogenic granuloma. J Clin Invest 1992, 90 (5), 1952–1957. DOI: 10.1172/jci116073.

(38) Karamanos, N. K.; Theocharis, A. D.; Piperigkou, Z.; Manou, D.; Passi, A.; Skandalis, S. S.; Vynios, D. H.; Orian-Rousseau, V.; Ricard-Blum, S.; Schmelzer, C. E. H.; et al. A guide to the composition and functions of the extracellular matrix. FEBS J 2021, 288 (24), 6850–6912. DOI: 10.1111/febs.15776.

(39) Patterson, J.; Hubbell, J. A. Enhanced proteolytic degradation of molecularly engineered PEG hydrogels in response to MMP-1 and MMP-2. Biomaterials 2010, 31 (30), 7836–7845. DOI: 10.1016/j.biomaterials.2010.06.061.

(40) Turk, B. E.; Huang, L. L.; Piro, E. T.; Cantley, L. C. Determination of protease cleavage site motifs using mixture-based oriented peptide libraries. Nat Biotechnol 2001, 19 (7), 661–667. DOI: 10.1038/90273.

(41) Xu, X.; Gurski, L. A.; Zhang, C.; Harrington, D. A.; Farach-Carson, M. C.; Jia, X. Recreating the tumor microenvironment in a bilayer, hyaluronic acid hydrogel construct for the growth of prostate cancer spheroids. Biomaterials 2012, 33 (35), 9049–9060. DOI: 10.1016/j.biomaterials.2012.08.061.

(42) Xu, X.; Sabanayagam, C. R.; Harrington, D. A.; Farach-Carson, M. C.; Jia, X. A hydrogel-based tumor model for the evaluation of nanoparticle-based cancer therapeutics. Biomaterials 2014, 35 (10), 3319–3330. DOI: 10.1016/j.biomaterials.2013.12.080.

(43) Hao, Y.; Zerdoum, A. B.; Stuffer, A. J.; Rajasekaran, A. K.; Jia, X. Biomimetic Hydrogels Incorporating Polymeric Cell-Adhesive Peptide To Promote the 3D Assembly of Tumoroids. Biomacromolecules 2016, 17 (11), 3750–3760. DOI: 10.1021/acs.biomac.6b01266.

(44) Lintz, M.; Muñoz, A.; Reinhart-King, C. A. The Mechanics of Single Cell and Collective Migration of Tumor Cells. J Biomech Eng 2017, 139 (2), 0210051–0210059. DOI: 10.1115/1.4035121.

(45) Cheung, K. J.; Padmanaban, V.; Silvestri, V.; Schipper, K.; Cohen, J. D.; Fairchild, A. N.; Gorin, M. A.; Verdone, J. E.; Pienta, K. J.; Bader, J. S.;, et al. Polyclonal breast cancer metastases arise from collective dissemination of keratin 14-expressing tumor cell clusters. Proc Natl Acad Sci U S A 2016, 113 (7), E854–863. DOI: 10.1073/pnas.1508541113.

(46) Aceto, N.; Bardia, A.; Miyamoto, D. T.; Donaldson, M. C.; Wittner, B. S.; Spencer, J. A.; Yu, M.; Pely, A.; Engstrom, A.; Zhu, H.;, et al. Circulating tumor cell clusters are oligoclonal precursors of breast cancer metastasis. Cell 2014, 158 (5), 1110–1122. DOI: 10.1016/j.cell.2014.07.013.

(47) Panková, K.; Rösel, D.; Novotný, M.; Brábek, J. The molecular mechanisms of transition between mesenchymal and amoeboid invasiveness in tumor cells. Cell Mol Life Sci 2010, 67 (1), 63–71. DOI: 10.1007/s00018-009-0132-1.

(48) Qin, G.; Luo, M.; Chen, J.; Dang, Y.; Chen, G.; Li, L.; Zeng, J.; Lu, Y.; Yang, J. Reciprocal activation between MMP-8 and TGF-β1 stimulates EMT and malignant progression of hepatocellular carcinoma. Cancer Lett 2016, 374 (1), 85–95. DOI: 10.1016/j.canlet.2016.02.001.

(49) Taubenberger, A. V.; Girardo, S.; Träber, N.; Fischer-Friedrich, E.; Kräter, M.; Wagner, K.; Kurth, T.; Richter, I.; Haller, B.; Binner, M.;, et al. 3D Microenvironment Stiffness Regulates Tumor Spheroid Growth and Mechanics via p21 and ROCK. Adv Biosyst 2019, 3 (9), e1900128. DOI: 10.1002/adbi.201900128.

(50) Yamada, K. M.; Sixt, M. Mechanisms of 3D cell migration. Nat Rev Mol Cell Biol 2019, 20 (12), 738–752. DOI: 10.1038/s41580-019-0172-9.

(51) Xia, P.; Gütl, D.; Zheden, V.; Heisenberg, C. P. Lateral Inhibition in Cell Specification Mediated by Mechanical Signals Modulating TAZ Activity. Cell 2019, 176 (6), 1379–1392.e1314. DOI: 10.1016/j.cell.2019.01.019.

(52) Shi, X.; Yang, J.; Deng, S.; Xu, H.; Wu, D.; Zeng, Q.; Wang, S.; Hu, T.; Wu, F.; Zhou, H. TGF-β signaling in the tumor metabolic microenvironment and targeted therapies. J Hematol Oncol 2022, 15 (1), 135. DOI: 10.1186/s13045-022-01349-6.

(53) Derynck, R.; Turley, S. J.; Akhurst, R. J. TGFβ biology in cancer progression and immunotherapy. Nat Rev Clin Oncol 2021, 18 (1), 9–34. DOI: 10.1038/s41571-020-0403-1.

(54) Zarzynska, J. M. Two faces of TGF-beta1 in breast cancer. Mediators Inflamm 2014, 2014, 141747. DOI: 10.1155/2014/141747.

(55) Meng, X.; Vander Ark, A.; Lee, P.; Hostetter, G.; Bhowmick, N. A.; Matrisian, L. M.; Williams, B. O.; Miranti, C. K.; Li, X. Myeloid-specific TGF-β signaling in bone promotes basic-FGF and breast cancer bone metastasis. Oncogene 2016, 35 (18), 2370–2378. DOI: 10.1038/onc.2015.297.

(56) Zhou, H.; Wu, G.; Ma, X.; Xiao, J.; Yu, G.; Yang, C.; Xu, N.; Zhang, B.; Zhou, J.; Ye, Z.;, et al. Attenuation of TGFBR2 expression and tumour progression in prostate cancer involve diverse hypoxia-regulated pathways. J Exp Clin Cancer Res 2018, 37 (1), 89. DOI: 10.1186/s13046-018-0764-9.

(57) Pang, Y.; Gara, S. K.; Achyut, B. R.; Li, Z.; Yan, H. H.; Day, C. P.; Weiss, J. M.; Trinchieri, G.; Morris, J. C.; Yang, L. TGF-β signaling in myeloid cells is required for tumor metastasis. Cancer Discov 2013, 3 (8), 936–951. DOI: 10.1158/2159-8290.Cd-12-0527.

(58) Zhang, Y. E. Non-Smad Signaling Pathways of the TGF-β Family. Cold Spring Harb Perspect Biol 2017, 9 (2). DOI: 10.1101/cshperspect.a022129.

(59) Wang, D.; Peng, L.; Hua, L.; Li, J.; Liu, Y.; Zhou, Y. Mapk14 is a Prognostic Biomarker and Correlates with the Clinicopathological Features and Immune Infiltration of Colorectal Cancer. Front Cell Dev Biol 2022, 10, 817800. DOI: 10.3389/fcell.2022.817800.

(60) Lee, H. W.; Park, Y. M.; Lee, S. J.; Cho, H. J.; Kim, D. H.; Lee, J. I.; Kang, M. S.; Seol, H. J.; Shim, Y. M.; Nam, D. H.;, et al. Alpha-smooth muscle actin (ACTA2) is required for metastatic potential of human lung adenocarcinoma. Clin Cancer Res 2013, 19 (21), 5879–5889. DOI: 10.1158/1078-0432.Ccr-13-1181.

(61) Jeon, M.; You, D.; Bae, S. Y.; Kim, S. W.; Nam, S. J.; Kim, H. H.; Kim, S.; Lee, J. E. Dimerization of EGFR and HER2 induces breast cancer cell motility through STAT1-dependent ACTA2 induction. Oncotarget 2017, 8 (31), 50570–50581. DOI: 10.18632/oncotarget.10843.

(62) Malekan, M.; Nezamabadi, S. S.; Samami, E.; Mohebalizadeh, M.; Saghazadeh, A.; Rezaei, N. BDNF and its signaling in cancer. J Cancer Res Clin Oncol 2023, 149 (6), 2621–2636. DOI: 10.1007/s00432-022-04365-8.

(63) Dasgupta, A.; Gibbard, D. F.; Schmitt, R. E.; Arneson-Wissink, P. C.; Ducharme, A. M.; Bruinsma, E. S.; Hawse, J. R.; Jatoi, A.; Doles, J. D. A TGF-β/KLF10 signaling axis regulates atrophy-associated genes to induce muscle wasting in pancreatic cancer. Proc Natl Acad Sci U S A 2023, 120 (34), e2215095120. DOI: 10.1073/pnas.2215095120.

(64) Jechlinger, M.; Sommer, A.; Moriggl, R.; Seither, P.; Kraut, N.; Capodiecci, P.; Donovan, M.; Cordon-Cardo, C.; Beug, H.; Grünert, S. Autocrine PDGFR signaling promotes mammary cancer metastasis. J Clin Invest 2006, 116 (6), 1561–1570. DOI: 10.1172/jci24652.

(65) Pal, S. K.; Nguyen, C. T.; Morita, K. I.; Miki, Y.; Kayamori, K.; Yamaguchi, A.; Sakamoto, K. THBS1 is induced by TGFB1 in the cancer stroma and promotes invasion of oral squamous cell carcinoma. J Oral Pathol Med 2016, 45 (10), 730–739. DOI: 10.1111/jop.12430.

(66) Roberts, E.; Cossigny, D. A.; Quan, G. M. The role of vascular endothelial growth factor in metastatic prostate cancer to the skeleton. Prostate Cancer 2013, 2013, 418340. DOI: 10.1155/2013/418340.

(67) Kopsida, M.; Liu, N.; Kotti, A.; Wang, J.; Jensen, L.; Jothimani, G.; Hildesjo, C.; Haapaniemi, S.; Zhong, W.; Pathak, S.;, et al. RhoB expression associated with chemotherapy response and prognosis in colorectal cancer. Cancer Cell Int 2024, 24 (1), 75. DOI: 10.1186/s12935-024-03236-1.

(68) Pickup, M.; Novitskiy, S.; Moses, H. L. The roles of TGFβ in the tumour microenvironment. Nat Rev Cancer 2013, 13 (11), 788–799. DOI: 10.1038/nrc3603.

(69) Goto, K.; Kamiya, Y.; Imamura, T.; Miyazono, K.; Miyazawa, K. Selective inhibitory effects of Smad6 on bone morphogenetic protein type I receptors. J Biol Chem 2007, 282 (28), 20603–20611. DOI: 10.1074/jbc.M702100200.

(70) Liu, B.; Mao, N. Smad5: signaling roles in hematopoiesis and osteogenesis. Int J Biochem Cell Biol 2004, 36 (5), 766–770. DOI: 10.1016/s1357-2725(03)00250-4.

(71) Zhang, Y.; Narayanan, S. P.; Mannan, R.; Raskind, G.; Wang, X.; Vats, P.; Su, F.; Hosseini, N.; Cao, X.; Kumar-Sinha, C.;, et al. Single-cell analyses of renal cell cancers reveal insights into tumor microenvironment, cell of origin, and therapy response. Proc Natl Acad Sci U S A 2021, 118 (24). DOI: 10.1073/pnas.2103240118.

(72) Hyakusoku, H.; Sano, D.; Takahashi, H.; Hatano, T.; Isono, Y.; Shimada, S.; Ito, Y.; Myers, J. N.; Oridate, N. JunB promotes cell invasion, migration and distant metastasis of head and neck squamous cell carcinoma. J Exp Clin Cancer Res 2016, 35, 6. DOI: 10.1186/s13046-016-0284-4.

(73) Lasorella, A.; Benezra, R.; Iavarone, A. The ID proteins: master regulators of cancer stem cells and tumour aggressiveness. Nat Rev Cancer 2014, 14 (2), 77–91. DOI: 10.1038/nrc3638.

(74) Strong, N.; Millena, A. C.; Walker, L.; Chaudhary, J.; Khan, S. A. Inhibitor of differentiation 1 (Id1) and Id3 proteins play different roles in TGFβ effects on cell proliferation and migration in prostate cancer cells. Prostate 2013, 73 (6), 624–633. DOI: 10.1002/pros.22603.

(75) Kong, H. J.; Kwon, E. J.; Kwon, O. S.; Lee, H.; Choi, J. Y.; Kim, Y. J.; Kim, W.; Cha, H. J. Crosstalk between YAP and TGFβ regulates SERPINE1 expression in mesenchymal lung cancer cells. Int J Oncol 2021, 58 (1), 111–121. DOI: 10.3892/ijo.2020.5153.

(76) Inman, G. J.; Nicolás, F. J.; Hill, C. S. Nucleocytoplasmic shuttling of Smads 2, 3, and 4 permits sensing of TGF-beta receptor activity. Mol Cell 2002, 10 (2), 283–294. DOI: 10.1016/s1097-2765(02)00585-3.

(77) Ao, M.; Williams, K.; Bhowmick, N. A.; Hayward, S. W. Transforming growth factor-beta promotes invasion in tumorigenic but not in nontumorigenic human prostatic epithelial cells. Cancer Res 2006, 66 (16), 8007–8016. DOI: 10.1158/0008-5472.Can-05-4451.

(78) Katsuno, Y.; Meyer, D. S.; Zhang, Z.; Shokat, K. M.; Akhurst, R. J.; Miyazono, K.; Derynck, R. Chronic TGF-β exposure drives stabilized EMT, tumor stemness, and cancer drug resistance with vulnerability to bitopic mTOR inhibition. Sci Signal 2019, 12 (570). DOI: 10.1126/scisignal.aau8544.

(79) Wang, Y.; Liu, J.; Ying, X.; Lin, P. C.; Zhou, B. P. Twist-mediated Epithelial-mesenchymal Transition Promotes Breast Tumor Cell Invasion via Inhibition of Hippo Pathway. Sci Rep 2016, 6, 24606. DOI: 10.1038/srep24606.

(80) Lamouille, S.; Xu, J.; Derynck, R. Molecular mechanisms of epithelial-mesenchymal transition. Nat Rev Mol Cell Biol 2014, 15 (3), 178–196. DOI: 10.1038/nrm3758.

(81) Hanada, S.; Harada, M.; Kumemura, H.; Omary, M. B.; Kawaguchi, T.; Taniguchi, E.; Koga, H.; Yoshida, T.; Maeyama, M.; Baba, S.;, et al. Keratin-containing inclusions affect cell morphology and distribution of cytosolic cellular components. Exp Cell Res 2005, 304 (2), 471–482. DOI: 10.1016/j.yexcr.2004.12.009.

(82) Jung, H.; Kim, B.; Moon, B. I.; Oh, E. S. Cytokeratin 18 is necessary for initiation of TGF- β1-induced epithelial-mesenchymal transition in breast epithelial cells. Mol Cell Biochem 2016, 423 (1-2), 21–28. DOI: 10.1007/s11010-016-2818-7.

(83) Munger, J. S.; Harpel, J. G.; Giancotti, F. G.; Rifkin, D. B. Interactions between growth factors and integrins: latent forms of transforming growth factor-beta are ligands for the integrin alphavbeta1. Mol Biol Cell 1998, 9 (9), 2627–2638. DOI: 10.1091/mbc.9.9.2627.

(84) Principe, D. R.; Doll, J. A.; Bauer, J.; Jung, B.; Munshi, H. G.; Bartholin, L.; Pasche, B.; Lee, C.; Grippo, P. J. TGF-β: duality of function between tumor prevention and carcinogenesis. J Natl Cancer Inst 2014, 106 (2), djt369. DOI: 10.1093/jnci/djt369.

