## Supplementary Information for "TGFβ1 and RGD Cooperatively Regulate SMAD 2/3 Mediated Oncogenic Effects in Prostate Cancer Cells in Bioorthogonally Constructed Hydrogels"

*Corresponding author:

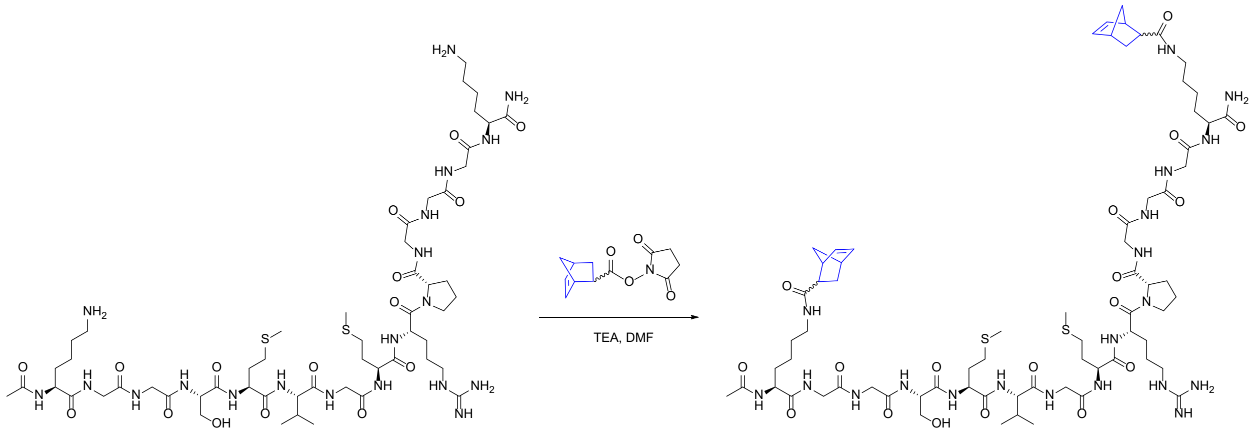


**Fig. S1.** Synthetic of Nb-conjugated non-degradable peptide crosslinker (SMV-bisNb).

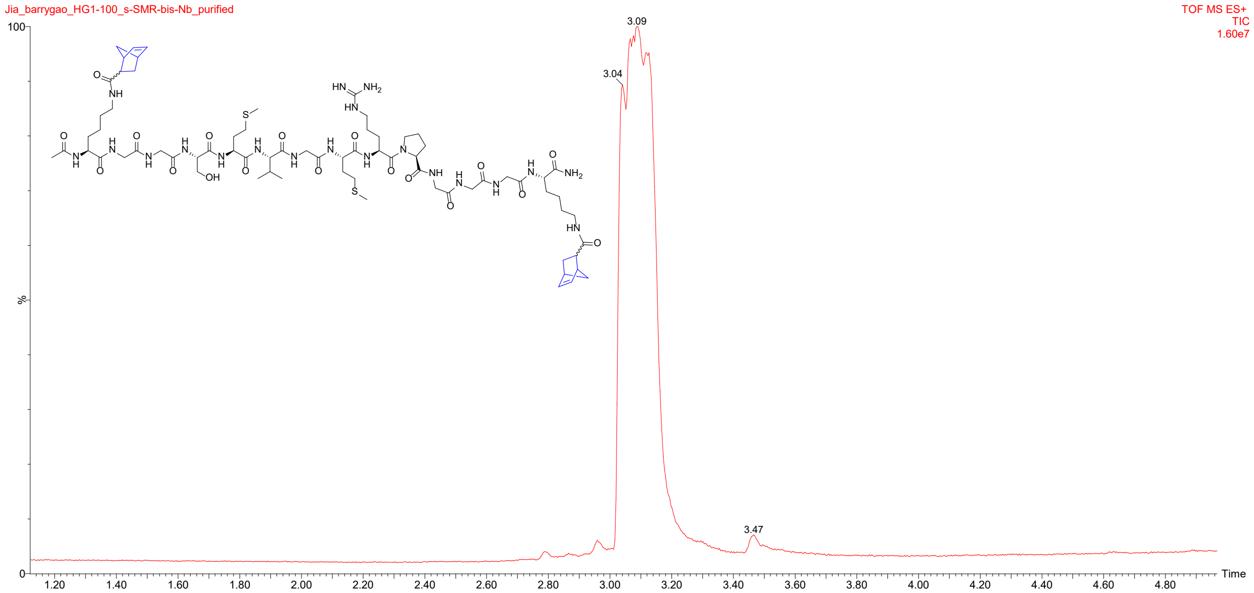


**Fig. S2.** UPLC-MS trace of purified SMV-bisNb crosslinker. Peak split was due to endo-exo isomerism of Nb.

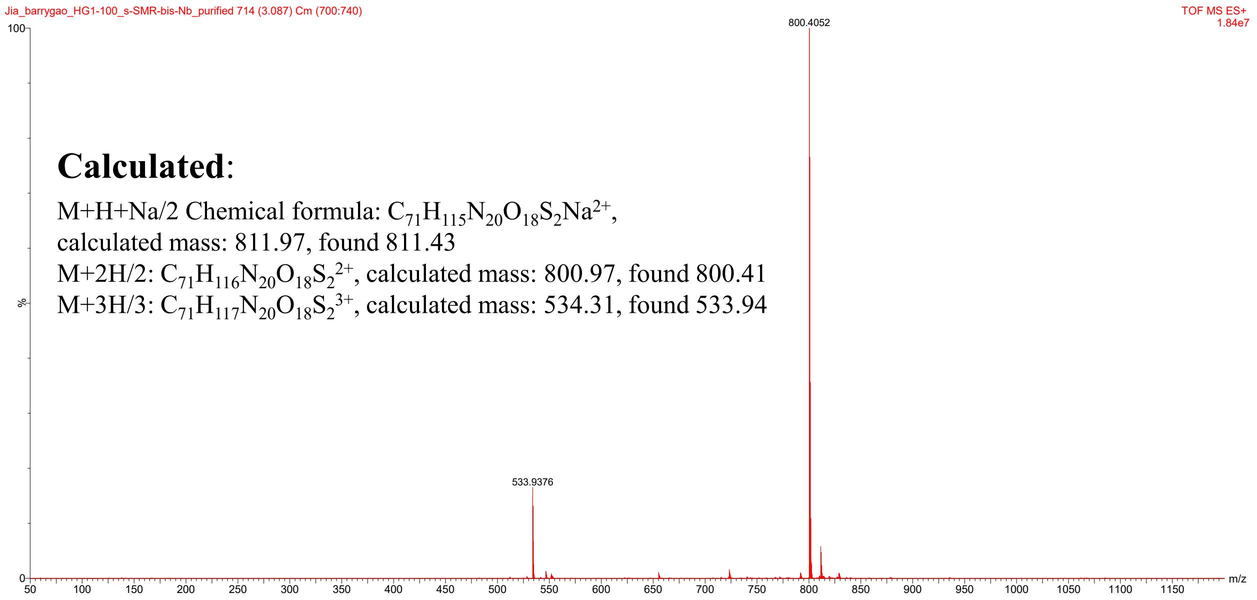


**Fig. S3.** UPLC-MS spectrum of purified SMV-bisNb crosslinker.

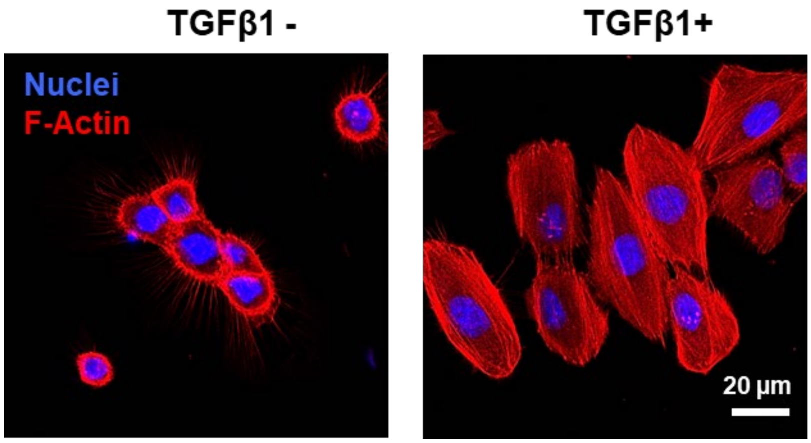


**Fig. S4.** Representative confocal images of DU145 cells cultured on 2D with or without TGFβ1 for 24 h. Cell nucleus was stained by DAPI (blue) and F-actin was stained by Alexa Fluor™ 568 phalloidin (red). Scale bar: 20 μm.

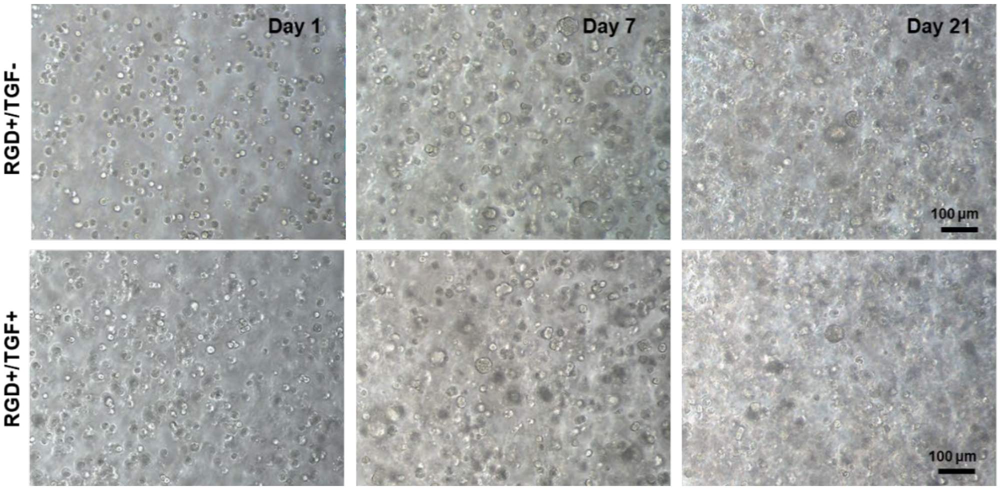


**Fig. S5.** Bright-field images of cellular constructs on days 1, 7, and 21. DU145 cells were encapsulated in non-MMP-degradable (crosslinker SMV-bisNb) with 1.0 mM RGD, with (10 ng/ml, RGD+/TGF+) or without (RGD+/TGF-) TGFβ1. TGFβ1 was introduced on day 3. Scale bar: 100 μm.

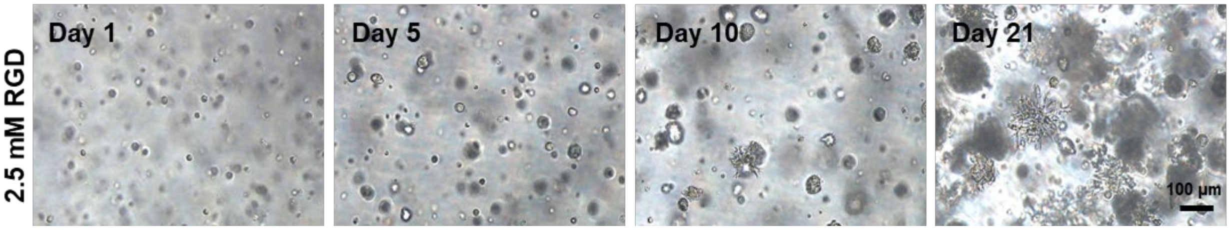


**Fig. S6.** Bright field images of DU145 cells maintained in MMP-degradable HA gels with 2.5 mM RGD without TGFβ1 for 21 days. Scale bar: 100 μm.

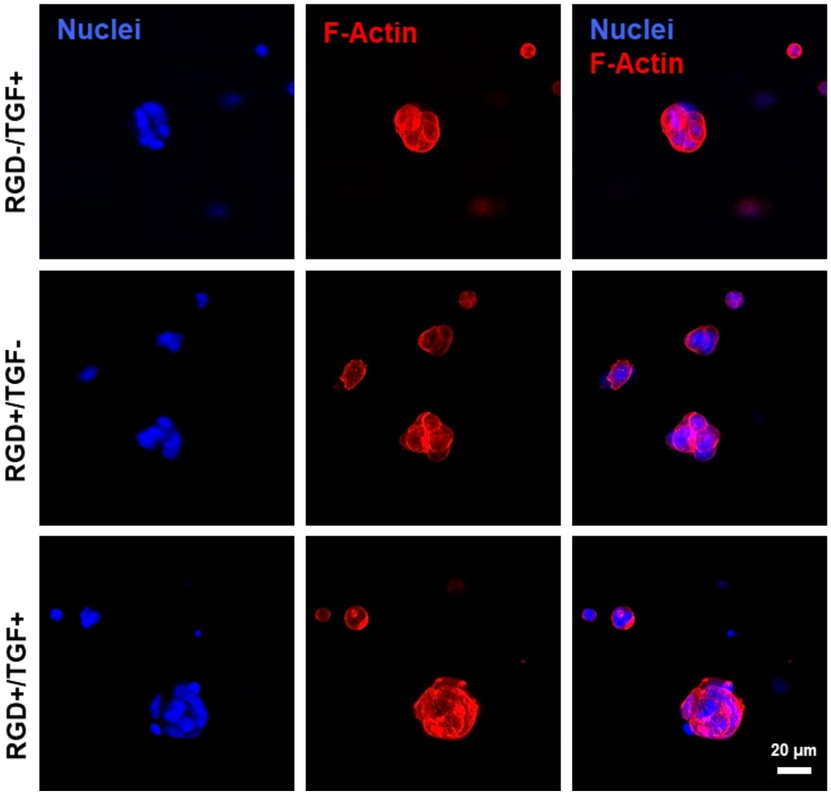


**Fig. S7.** Representative confocal images of RGD-/TGF+, RGD+/TGF- and RGD+/TGF+ constructs on day 3. Nuclei: blue, F-actin: red. Scale bar: 20 µm.

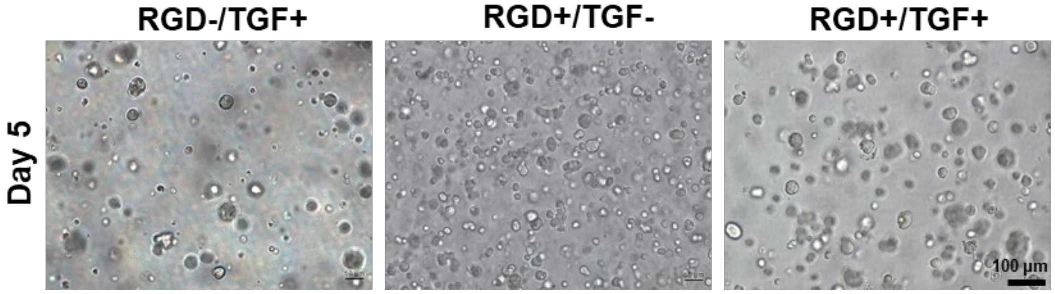


**Fig. S8.** Bright field images of DU145 cells maintained in MMP-degradable HA gels with or without RGD (1.0 mM) and TGFβ1 (10 ng/mL) on day 5 of culture. Scale bar: 100 µm.

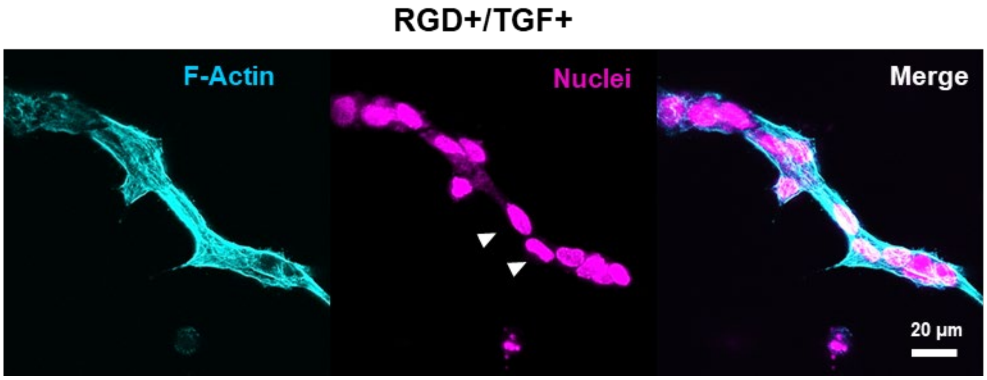


**Fig. S9.** Representative confocal images of DU145 cell on day 21 in RGD+/TGF+ cultures. Cell nuclei: magenta, F-actin: cyan. White arrowheads point to disseminated single cells. Scale bar: 20 µm.

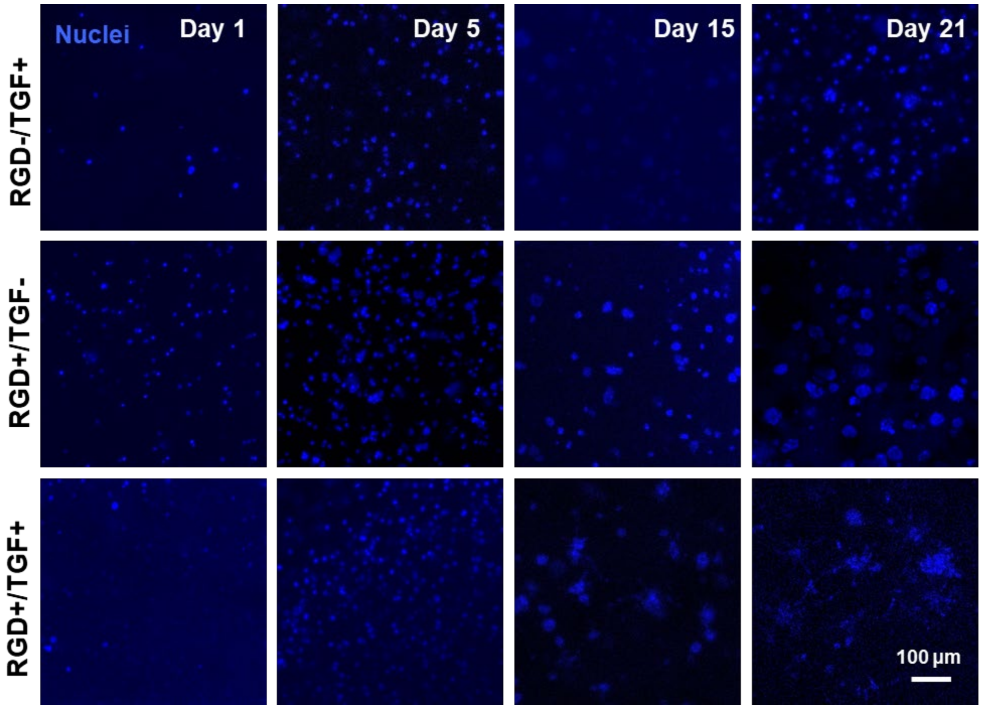


**Fig. S10.** Representative confocal images of 3D cultures stained with Hoechst for live/dead quantification. Scale bar: 100 µm.

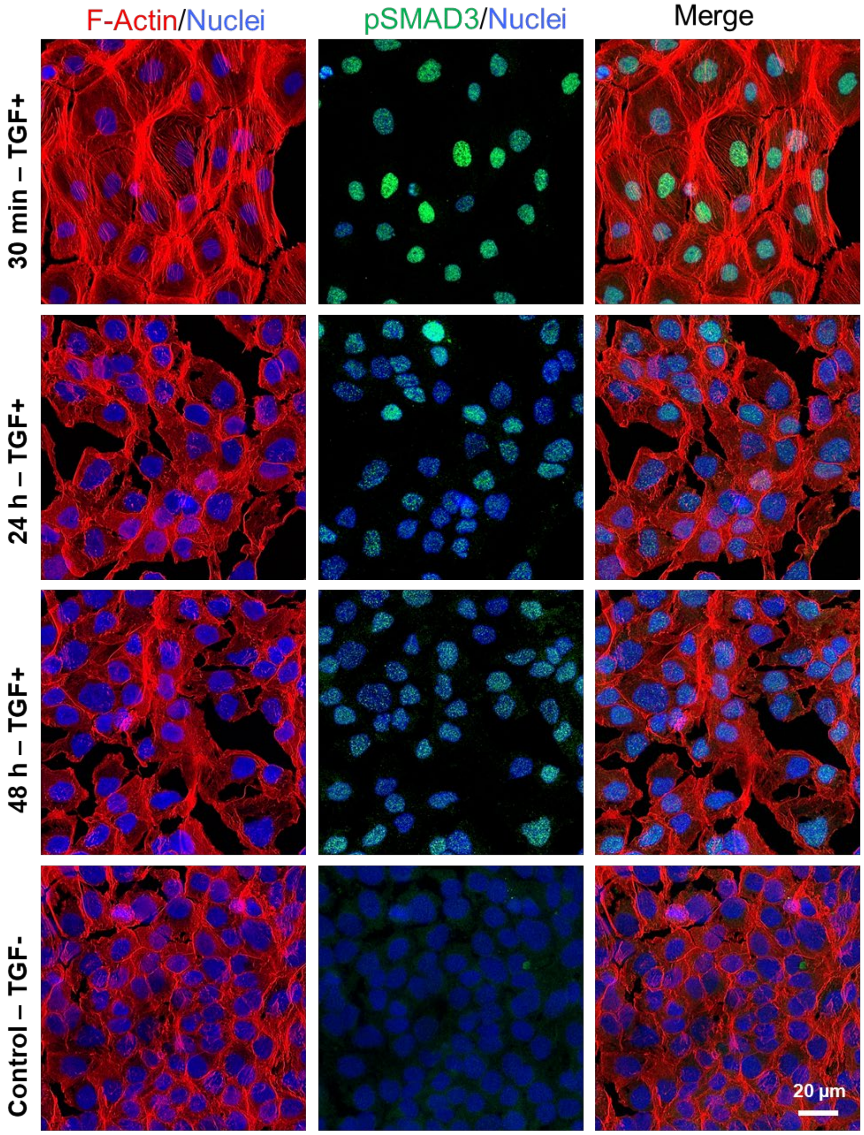


**Fig. S11.** Representative confocal images showing nuclear localization of pSMAD3 protein in DU145 cells cultured on 2D with or without TGFβ1 treatment (control). Cells were exposed to TGFβ1 for 30 min, 24 h, and 48 h before the analysis was performed. pSMAD3: green, Nuclei: blue, F-actin: red. Scale bar: 20 µm.

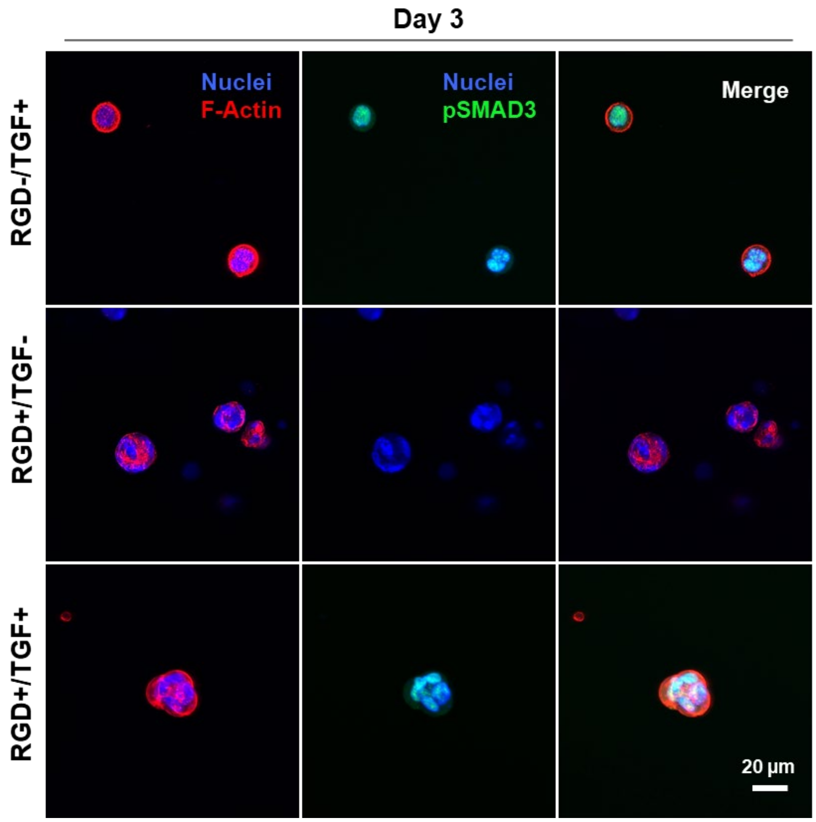


**Fig. S12**. Representative confocal images showing nuclear localization of pSMAD3 protein in DU145 cells cultured in 3D with or without TGFβ1 treatment. Cells were exposed to TGFβ1 for 30 min on day 3 before the analysis was performed. pSMAD3: green, Nuclei: blue, F-actin: red. Scale bar: 20 µm.

**Table S1**. The RT2 PCR array contains TGFβ1 target genes that belong to the differentiation and development functional category.

| Gene Full Form | Gene Abbreviation |
|---|---|
| Angiotensinogen (serpin peptidase inhibitor, clade A, member 8) | AGT |
| Bromodomain containing 2 | BRD2 |
| DnaJ (Hsp40) homolog, subfamily A, member 1 | DNAJA1 |
| Epithelial membrane protein 1 | EMP1 |
| Endoglin | ENG |
| EPH receptor B2 | EPHB2 |
| Fibronectin 1 | FN1 |
| FBJ murine osteosarcoma viral oncogene homolog | FOS |
| Interferon-related developmental regulator 1 | IFRD1 |
| Mitogen-activated protein kinase 14 | MAPK14 |
| Matrix metallopeptidase 2 (gelatinase A, 72kDa gelatinase, 72kDa type IV collagenase) | MMP2 |
| Nuclear factor of kappa light polypeptide gene enhancer in B-cells inhibitor, alpha | NFKBIA |
| Snail homolog 1 (Drosophila) | SNAI1 |
| Transforming growth factor, beta 2 | TGFB2 |
| Transforming growth factor, beta receptor II (70/80kDa) | TGFBR2 |

**Table S2.** The RT2 PCR array contains TGFβ1 target genes that belong to cell invasion and migration functional category.

| Gene Full Form | Gene Abbreviation |
|---|---|
| Actin, alpha 2, smooth muscle, aorta | ACTA2 |
| BCL2-like 1 | BCL2L1 |
| Brain-derived neurotrophic factor | BDNF |
| Cyclin-dependent kinase inhibitor 1B (p27, Kip1) | CDKN1B |
| Furin (paired basic amino acid cleaving enzyme) | FURIN |
| Heme oxygenase (decycling) 1 | HMOX1 |
| Interleukin 10 | IL10 |
| Kruppel-like factor 10 | KLF10 |
| Msh homeobox 2 | MSX2 |
| V-myc myelocytomatosis viral oncogene homolog (avian) | MYC |
| Platelet-derived growth factor alpha polypeptide | PDGFA |
| Plasminogen | PLG |
| Prostaglandin-endoperoxide synthase 2 (prostaglandin G/H synthase and cyclooxygenase) | PTGS2 |
| Parathyroid hormone-like hormone | PTHLH |
| PTK2 protein tyrosine kinase 2 | PTK2 |
| PTK2B protein tyrosine kinase 2 beta | PTK2B |
| Serpin peptidase inhibitor, clade E (nexin, plasminogen activator inhibitor type 1), member 1 | SERPINE1 |
| Sonic hedgehog | SHH |
| SRY (sex determining region Y)-box 4 | SOX4 |
| Thrombospondin 1 | THBS1 |

**Table S3**. The RT2 PCR array contains TGFβ1 target genes that belong to cell cycle regulation functional category.

| Gene Full Form | Gene Abbreviation |
|---|---|
| Activin A receptor, type I | ACVR1 |
| Cell division cycle 6 homolog (S. cerevisiae) | CDC6 |
| Ras homolog gene family, member B | RHOB |

**Table S4**. The RT2 PCR array contains TGFβ1 target genes that belong to anti-apoptotic/angiogenesis functional category.

| Gene Full Form | Gene Abbreviation |
|---|---|
| CCAAT/enhancer binding protein (C/EBP), beta | CEBPB |
| Crystallin, alpha B | CRYAB |
| Vascular endothelial growth factor A | VEGFA |

**Table S5.** The RT2 PCR array contains TGFβ1 target genes that belong to apoptosis functional category.

| Gene Full Form | Gene Abbreviation |
|---|---|
| Aryl hydrocarbon receptor interacting protein-like 1 | AIPL1 |
| Growth arrest and DNA-damage-inducible, beta | GADD45B |
| Homocysteine-inducible, endoplasmic reticulum stress-inducible, ubiquitin-like domain member 1 | HERPUD1 |
| Mitogen-activated protein kinase kinase kinase 7 | MAP3K7 |
| Mitogen-activated protein kinase 8 | MAPK8 |
| RAD21 homolog (S. pombe) | RAD21 |
| RING1 and YY1 binding protein | RYBP |
| S100 calcium binding protein A8 | S100A8 |
| Tumor necrosis factor (ligand) superfamily, member 10 | TNFSF10 |

**Table S6**. The RT2 PCR array contains TGFβ1 target genes that belong to the signal transduction functional category

| Gene Full Form | Gene Abbreviation |
|---|---|
| Thioredoxin interacting protein | TXNIP |
| Activin A receptor type II-like 1 | ACVRL1 |
| CREB binding protein | CREBBP |
| Catenin (cadherin-associated protein), beta 1, 88kDa | CTNNB1 |
| E2F transcription factor 4, p107/p130-binding | E2F4 |
| E1A binding protein p300 | EP300 |
| GLI family zinc finger 2 | GLI2 |
| General transcription factor IIi | GTF2I |
| Hairy and enhancer of split 1, (Drosophila) | HES1 |
| Hairy/enhancer-of-split related with YRPW motif 1 | HEY1 |
| Inhibitor of DNA binding 1, dominant negative helix-loop-helix protein | ID1 |
| Inhibitor of DNA binding 2, dominant negative helix-loop-helix protein | ID2 |
| Inhibitor of DNA binding 3, dominant negative helix-loop-helix protein | ID3 |
| Retinoblastoma-like 1 (p107) | RBL1 |
| Ras homolog gene family, member A | RHOA |
| SMAD family member 1 | SMAD1 |
| SMAD family member 3 | SMAD3 |
| SMAD family member 5 | SMAD5 |
| SMAD family member 6 | SMAD6 |
| Sp1 transcription factor | SP1 |

**Table S7**. The RT2 PCR array contains TGFβ1 target genes that belong to transcription factors functional category.

| **Gene Full Form** | **Gene Abbreviation** |
|---|---|
| Androgen receptor | AR |
| Activating transcription factor 3 | ATF3 |
| Activating transcription factor 4 (tax-responsive enhancer element B67) | ATF4 |
| BTB and CNC homology 1, basic leucine zipper transcription factor 1 | BACH1 |
| Basic helix-loop-helix family, member e40 | BHLHE40 |
| CAMP responsive element binding protein 1 | CREB1 |
| Methyl-CpG binding domain protein 1 | MBD1 |
| Myogenic differentiation 1 | MYOD1 |
| Nuclear factor I/B | NFIB |
| Notch 1 | NOTCH1 |
| Peroxisome proliferator-activated receptor alpha | PPARA |
| Retinoic acid receptor, alpha | RARA |
| Runt-related transcription factor 1 | RUNX1 |
| Sterol regulatory element binding transcription factor 2 | SREBF2 |

**Table S8.** List of primers used for RT qPCR analyses.

| Gene Name | Gene symbols | Forward primer (5'-3') | Reverse primer (5'-3') |
|---|---|---|---|
| Jun-B | *JUNB* | ACGACTCATACACAGCTACGG | GCTCGGTTTCAGGAGTTTGTAGT |
| Vimentin | *VIM1* | TGTCCAAATCGATGTGGATGTTTC | TTGTACCATTCTTCTGCCTCCTG |
| E-cadherin | *CDH1* | CGA GAG CTA CAC GTT CAC GG | GGG TGT CGA GGG AAA AAT AGG |
| Twist Basic Helix-Loop-Helix Transcription Factor 1 | *TWIST1* | AGCAAGATTCAGACCCTCAAGCT | CCTGGTAGAGGAAGTCGATGTACCT |
| Matrix metalloproteinase-9 | *MMP9* | TGTACCGCTATGGTTACACTCG | GGCAGGGACAGTTGCTTCT |
| Membrane type-1 matrix metalloproteinase | *MT1-MMP* | GGCTACAGCAATATGGCTACC | GATGGCCGCTGAGAGTGAC |
| SMAD 2 | SMAD2 | CGTCCATCTTGCCATTCACG | CTCAAGCTCATCTAATCGTCCTG |
| SMAD 4 | SMAD4 | TCCCAACATTCCTGTGGCTTC | CTGCTGCTGTCCTGGCTGA |
| Interleukin 6 | *IL-6* | ACTCACCTCTTCAGAACGAATTG | CCATCTTTGGAAGGTTCAGGTTG |
| Ki-67 | *Ki-67* | TGGGTCTGTTATTGATGAGCC | CATCAGGGTCAGAAGAGAAGC |

**Table S9**. Antibodies and their conditions used for immunocytochemistry.

| **Target** | **Vendor** | **Clone** | **Catalog number** | **Host** | **Dilution (v/v)** |
|---|---|---|---|---|---|
| E-cadherin | Abcam | EP700Y | ab40772 | Rabbit | 1/50 |
| Vimentin | Genway Biotech | VMT-24 | GWB-BBB094 | Mouse | 1/200 |
| P-SMAD2/3 | Abcam | EP823Y | ab52903 | Rabbit | 1/50 |
| Jun-B | Invitrogen | 15H19L4 | 701702 | Rabbit | 1/50 |
| ZO-1 | Invitrogen | ZO1-1A12 | 740002M | Mouse | 1/50 |
| Keratin 18 | Invitrogen | DC10 | MA5-12104 | Mouse | 1/200 |
| Keratin 19 | Abcam | EP1580Y | ab52625 | rabbit | 1/200 |
